# A phloem-localized Arabidopsis metacaspase (AtMC3) improves drought tolerance

**DOI:** 10.1101/2022.11.09.515759

**Authors:** Eugenia Pitsili, Ricardo Rodriguez-Trevino, Nerea Ruiz-Solani, Fatih Demir, Elizabeth Kastanaki, Charlene Dambire, Roger de Pedro, Dominique Vercammen, Jose Salguero-Linares, Hardy Hall, Melissa Mantz, Martin Schuler, Hannele Tuominen, Frank Van Breusegem, Marc Valls, Sergi Munné-Bosch, Michael J. Holdsworth, Pitter F. Huesgen, Antia Rodriguez-Villalon, Nuria S. Coll

## Abstract

Increasing drought phenomena pose a serious threat to agricultural productivity. Although plants have multiple ways to respond to the complexity of drought stress, the underlying mechanisms of stress sensing and signalling remain unclear. The role of the vasculature, in particular the phloem, in facilitating inter-organ communication is critical. Here, we investigated the role of AtMC3, a phloem-specific member of the metacaspase family, in osmotic stress responses in *Arabidopsis thaliana*. Overexpression of *AtMC3* conferred drought tolerance by enhancing the differentiation of the metaphloem sieve elements and maintaining higher levels of vascular-mediated transportation, whilst plants lacking the protein showed an impaired response to drought and inability to respond effectively to the hormone abscisic acid. Analyses of the proteome in plants with altered AtMC3 levels revealed differential abundance of proteins related to osmotic stress. Overall, our data highlight the importance of AtMC3 and vascular plasticity in fine-tuning early drought responses at the whole plant level without affecting growth or yield.

## Introduction

Climate change is an unfolding threat that is becoming increasingly prevalent due to its huge impact in many aspects of life. Agricultural productivity is negatively affected by climate change and in combination with population growth and increasing food demand, it has become urgent for scientists to identify new ways to cope with its related challenges^1^. An increase in drought events is one of the most frequent and severe consequences of climate change, reducing crop yield and productivity by 10% every year^2, 3^. Plants, as sessile organisms, have evolved ways to endure water scarcity and periods of drought stress. Yet, the underlying tissue-specific mechanisms resulting in the adaptation of plant growth and architecture to limited water conditions are not completely understood. Only by improving our understanding about the contribution of each tissue in plant acclimation to such a complex stress as drought, we will be able to translate this knowledge to biotechnological and breeding solutions and improve crop performance in the field^4^.

When facing drought stress, plants arrest growth, close stomata and start accumulating osmoprotectant molecules to prevent excessive water loss and minimize cellular damage^5^. As a result of stomatal closure, photosynthesis is arrested, leading to a negative carbon balance and carbon deprivation which may cause mortality in the long run^6^. Many of these physiological responses are triggered by the synthesis and perception of the hormone abscisic acid (ABA), an essential orchestrator of abiotic stress resistance mechanisms that regulates water status to protect cell systems and induces the expression of a gene network including dehydration tolerant proteins^7, 8^. To coordinate an efficient response to water deprivation at the organismal level, plants rely on their vascular system connecting distantly separated organs. In higher plants, such as *Arabidopsis thaliana* (hereafter Arabidopsis), the vascular system comprises xylem and phloem tissues. While the main function of xylem is the root-to-shoot transport of water and nutrients, phloem distributes photoassimilates and growth regulators throughout the whole plant body. It is well accepted that drought stress triggers the translocation of root-synthesized ABA to the shoot through the xylem, while coordinating stomatal closure^9, 10^. Recent studies have demonstrated that drought stress promotes endodermal production of ABA^11–13^, which is then translocated to the xylem where it modulates the signaling cascades governing its specification and differentiation^14^. Most of these studies have focused on the modification of xylem architecture as a response to the stress, whereas in the case of phloem, knowledge is limited to the fact that drought-associated osmotic imbalance may promote the release of water from phloem cells and eventually collapse tissue transport. While loss of xylem conductivity is a hallmark of plant death, it has been proposed that the timing of phloem collapse could determine not only the survival but also the revival capacity of the plants after drought, making this tissue an important factor for forecasting the plant behavior under water scarcity^15^.

In Arabidopsis, two types of conductive elements constitute phloem tissue, protophloem and metaphloem sieve elements (PSE and MSE respectively), whose survival depends on the activity of neighboring companion cells (CCs) (Fig. 1A). Unlike metaphloem, the molecular mechanism underpinning protophloem specification and differentiation has been extensively studied during the last decade (^16, 17^). PSEs are the first cells to differentiate within the root meristem and they orchestrate the development of their neighboring tissues. The functional association of CC and PSEs provides developmental plasticity to root cells safeguarding the functionality of the phloem tissues^18, 19^. PSE early differentiation is required for the root meristematic unloading of sugars and hormones^20^. Since sugars constitute the energy source for root development, defects in protophloem differentiation are reflected in an impaired meristematic activity and, in turn, post-embryonic root growth^21, 22^. Less studied is the plasticity of the tissue in response to environmental stresses, although proteomic profiling of exudates from plants exposed to drought stress has revealed that the phloem transports water deficit-related signals^23^.

**Figure 1:**
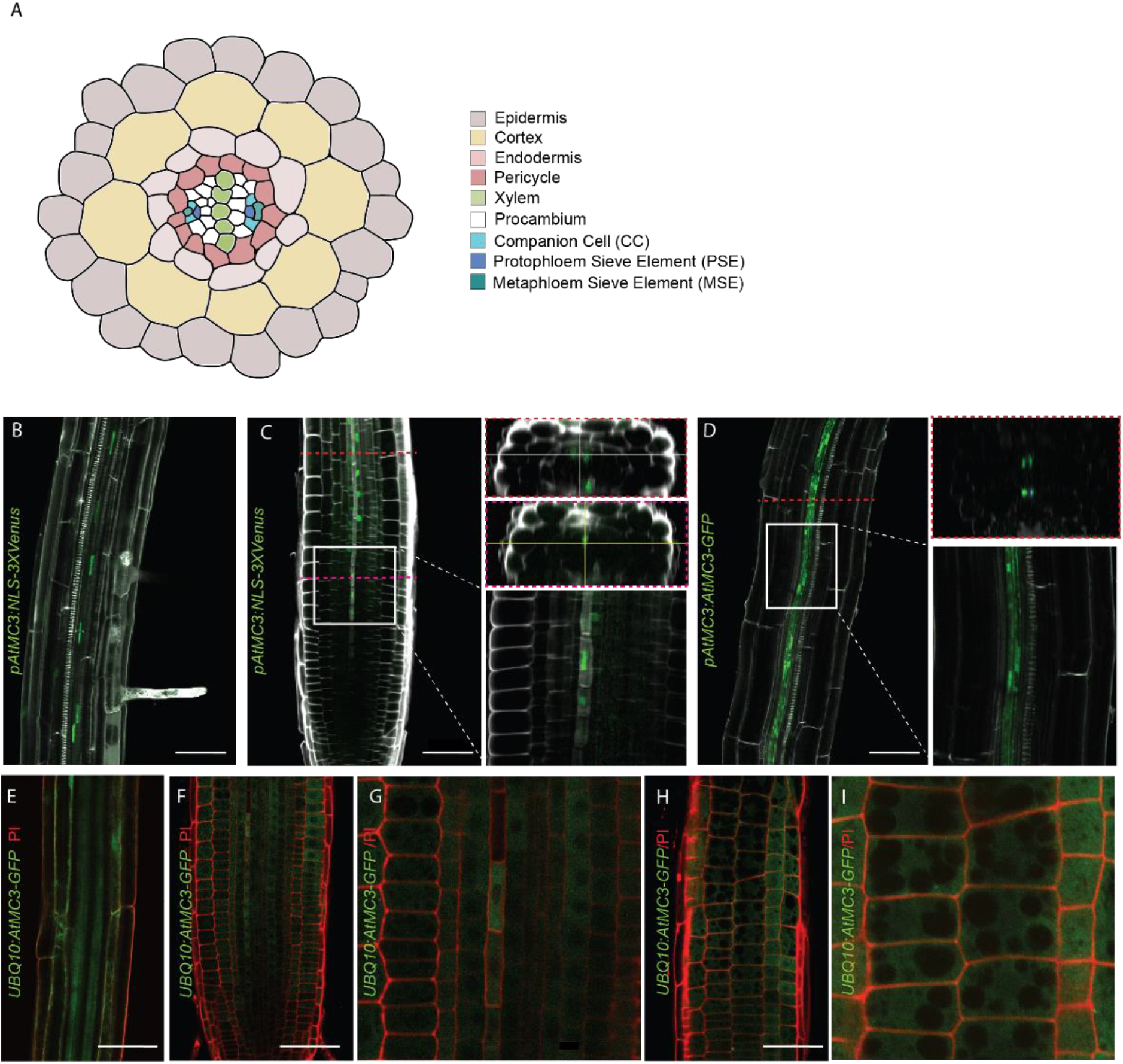
AtMC3 shows phloem-specific expression. (A) Schematic representation of a radial view of primary root in Arabidopsis thaliana with color-coded cell types. (B, C) Expression pattern of AtMC3 gene as revealed by pAtMC3: NLS-3xVenus construct with confocal microscopy in the root differentiation zone (B) and in the root meristem (C) of 6-day-old seedlings. Nuclear-localized mVENUS signals (green) are co-visualized with calcofluor staining (white) in the CCs of the root. (C) On the right lower panel is the magnification of the vascular tissue from the meristematic zone at the protophloem enucleation point. Transverse observations of the root are also present on the right upper panel. (D) Expression pattern of AtMC3 protein monitored with pAtMC3:AtMC3-GFP in the root differentiation zone in 7-day-old seedlings. Translational fusions with GFP signals (green) are co-visualized with propidium iodide (white). On the right lower panel is the magnification of the vascular tissue for the cytoplasmic companion cell visualization. Transverse observation of the root is also present on the right upper panel. (E-I) Expression pattern of AtMC3 monitored with pUBQ10:AtMC3-GFP in the root of 7-day-old seedlings. (E) Differentiation zone of the root, (F, G) meristematic zone focusing on the vasculature with magnification, (H, I) meristematic zone focusing on the epidermal cells with magnification. Translational fusions with GFP signals (green) are co-visualized with propidium iodide (red). Scale bar, 50 μm.

Metacaspases are a group of cysteine proteases that participate in various stress responses^24, 25^. This family of proteins are found in plants and lower eukaryotes and have some structural resemblance to the animal cell death regulators known as caspases, as they bear a catalytic domain comprising the p20 and p10 subunits. However, their mode of action is considerably different from caspases as they are calcium-dependent and cleave substrates after a lysine or an arginine^26–28^. In plants, metacaspases can be divided into type I and type II sub-groups. While Type I metacaspases have an N-terminal prodomain, type II lack the prodomain but harbour a linker region between their p20 and p10 catalytic subunits. The role of metacaspases has been mostly investigated in the context of stress responses and regulated cell death in Arabidopsis, which carries 9 metacaspase genes, three type I (AtMC1/AtMCAIa-AtMC3/AtMCAIc) and six type II (AtMC4/AtMCAIIa-AtMC9/AtMCAIIf)^29–33^.

Here, we show that the Arabidopsis type I metacaspase AtMC3/AtMCAIc displays specific expression in the phloem vascular tissue within the root. Alteration of AtMC3 levels leads to differential accumulation of stress-related proteins, indicating that AtMC3 is specifically involved in regulating osmotic stress responses. We report for the first time that a phloem-localized metacaspase contributes to drought tolerance by enhancing the differentiation of metaphloem vascular tissue and maintaining effective long-distance transmission of molecules under osmotic stress conditions. Additionally, we provide evidence demonstrating that reduced levels of AtMC3 affect the sensitivity of the plant to ABA.

## Results

### AtMC3 is a phloem-specific metacaspase

*AtMC3* expression is specifically detected in vascular tissues of all plant organs in young seedlings, as revealed by transgenic Arabidopsis lines carrying the *AtMC3* promoter fused to GUS (*pAtMC3:GUS*) (Fig. S1A-D) and as has been described before^19, 34^. Using mature plants, we observed that *AtMC3* expression was mostly restricted to the phloem, as shown by transverse cuts either in the stem or the hypocotyl (Fig. S1E-L). Specifically, in the phloem of the hypocotyl, the expression appears to be strongest in a small cell type that represents CCs (Fig S1G). To determine in more detail the cell specific expression of *AtMC3,* Arabidopsis reporter lines containing nuclear-localized fluorescent VENUS protein under the endogenous *AtMC3* promoter were generated (*pAtMC3:NLS-3xVENUS*). Confocal microscopy analysis of *pAtMC3:NLS-3xVENUS* roots confirmed the phloem pole specific expression of *AtMC3* (Fig. 1B, C). Initiation of *pAtMC3*-driven *NLS-3xVENUS* expression within the root meristem was observed in PSE differentiating cells prior to their enucleation. After completion of the PSE differentiation process, NLS-3xVENUS was detected in the CCs of the elongation zone (Fig. 1C). Additionally, *pAtMC3*-driven *NLS-3xVENUS* expression was also specifically detected at the CCs within the transition and from the differentiation zone onwards in the root (Fig. 1B).

To identify whether *AtMC3* expression in the phloem corresponded to the site of AtMC3 protein accumulation or transport through the vasculature, we generated transgenic lines stably expressing *AtMC3* fused to green fluorescent protein (GFP) under the control of *AtMC3* promoter (*pAtMC3:AtMC3-GFP*). Confocal analysis revealed that AtMC3-GFP specifically localizes in CCs in the differentiation zone and the signal could also be detected in the differentiating PSE in the meristematic zone (Fig. 1D, S2A, B). As expected, AtMC3 overexpression (*pUBQ10:AtMC3-GFP*) results in protein accumulation throughout the root, (Fig. 1E, F). Interestingly, GFP signal was not detected in enucleated SEs, suggesting that AtMC3 cannot be transferred through the vasculature (Fig. 1G). Subcellularly, *pAtMC3*-driven *AtMC3* expression appeared evenly distributed throughout the cytoplasm (Fig. 1C), which is in accordance with the localization of most of metacaspases studied^30–32^. Using the overexpression line to visualize the epidermal cells of the root meristem that are bigger in size, AtMC3-GFP fluorescent signal was not detected in the nucleus, nor in the vacuoles, suggesting that AtMC3 localizes in the cytosol (Fig. 1H, I). Mutation of the predicted catalytic cysteine to an alanine in AtMC3 (C230A) (Fig. 2A) did not affect protein localization, (Fig. S2C-F), but resulted in increased accumulation of the protein (Fig S2G), as previously observed for other metacaspases, such as AtMC1^35^.

**Figure 2:**
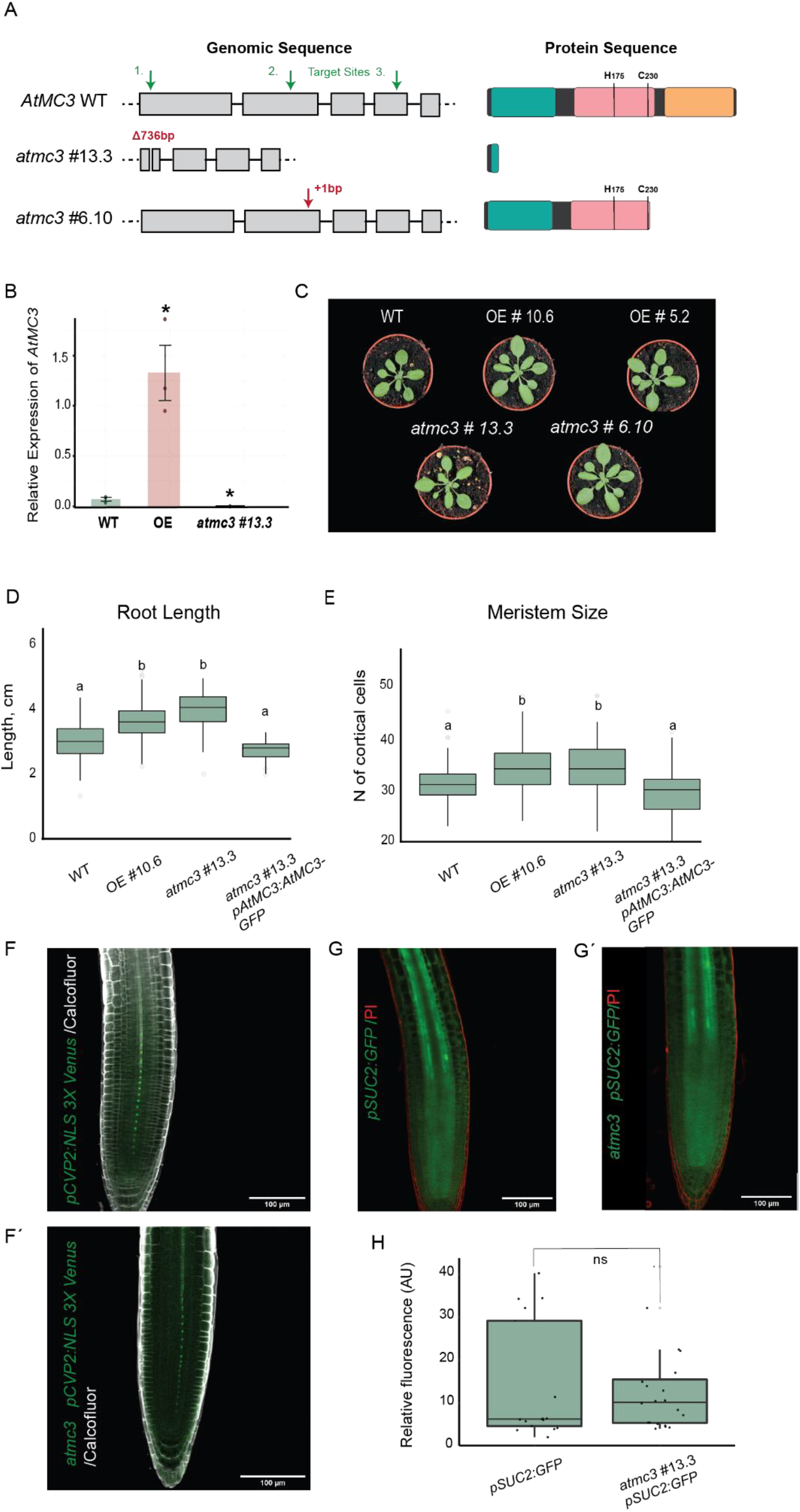
Characterization of AtMC3 CRISPR mutant. (A) Schematic representation of AtMC3 gene and protein structure of Col-0 WT and CRISPR mutant plants generated. Target sites chosen are indicated with green arrows and mutations generated are indicated with red text on top of the gene loci affected. The different grey boxes represent the exons of the gene sequence. In the protein structure, green colour is representing the metacaspase prodomain, pink and orange colours are indicating the catalytic p20 and p10 domains respectively. The predicted catalytic sites are indicated with H for Histidine in position 174 and C for Cysteine in position 230. (B) Expression analysis for the AtMC3 gene in WT, overexpressor and atmc3 #13.3 plants, in 7-day-old seedlings roots. Data are means (±SD) of three biological replicates. Significant differences from WT were determined by Student’s t-test: *P<0.05. (C) Rosette phenotypes of 3-week-old plants for WT, two independent overexpressor lines and two CRISPR mutant lines. (D) Root length of 7-day-old seedlings and (E) meristematic size of 6-day-old seedlings measured from WT, overexpressor, atmc3 #13.3mutant and complemented plants with the gene under the endogenous promoter. Different letters depict significant differences in a one-way ANOVA plus a Tukey’s HSD test. (F, F′) Representative pictures of confocal microscopy from root tips for observation of PSE nuclei in WT and atmc3 #13.3 mutant plants carrying pCVP2:NLS-3xVenus. (G, G′) Representative pictures from root tips of WT and atmc3 #13.3 mutant plants carrying pSUC2:GFP. (H) Quantification of unloaded cytosolic GFP of plants from (G, G′), (±SE). Fluorescence intensity was quantified over a rectangular area 100 μm above the QC; n≥12 roots.

### Altered expression of *AtMC3* enhances seedling root growth without affecting phloem formation

In order to define the role of AtMC3 within phloem tissues, CRISPR genome editing technology was employed to generate targeted mutations^36^. Homozygous lines were obtained each containing a different mutation in the sequence of *AtMC3* that resulted in C-terminal truncations in the proteins due to the frame alteration and generation of premature stop codons (Fig. 2A): *atmc3* #6.10 and #13.3. The latter presents a 763 bp deletion and was therefore selected for further functional analysis. *AtMC3* transcript levels demonstrated that *atmc3* #13.3 plants are null mutants, incapable of producing full-length transcripts, whereas transgenic lines overexpressing *AtMC3* show enhanced levels of expression compared to wild-type (WT) plants (Fig. 2B).

To determine whether AtMC3 is involved in plant growth or development, a detailed phenotypic characterization of the *atmc3* mutants and overexpressor lines was performed. No differences were observed at the adult stage in terms of rosette size, plant height or inflorescence tissues (Fig. 2C, S3A). However, at the seedling stage, both null mutant and overexpressing lines exhibited a larger root meristem size compared to WT plants, which translated into significantly longer roots (Fig. 2D, E). In addition, seedlings were analyzed for phloem deficit-related phenotypes such as lateral root formation and cotyledon vein pattern, but no differences were detected between the lines examined (Fig. S3C, D). Confocal microscopy analysis did not reveal any defects in protophloem continuity (Fig. S3B), an observation confirmed by the continuous nuclear expression of *GFP* driven by the protophloem-specific promoter *COTYLEDON VASCULAR PATTERN 2 (CVP2)* (Fig. 2F, F′). Furthermore, protophloem-mediated unloading in the root tip was not affected in the *atmc3* #13.3 mutant as revealed by the detection and quantification of *GFP* expressed under the specific companion cell promoter *SUCROSE TRASPORTER 2 (SUC2*) in root meristem cells (Fig. 2G, G′, H).

### *atmc3* null mutant displays reduced sensitivity to the stress hormone abscisic acid (ABA)

To assess the effects of altering the levels of AtMC3 in the plant and whether its putative protease function is relevant, we undertook a proteomic approach. Our initial idea was to perform an analysis comparing the degradomes of the *atmc3#13.3* mutant and *AtMC3* overexpressor to WT plants, aimed at identifying potential proteolytic substrates of AtMC3, as the protein is a predicted cysteine protease^37^. For this, we carried out High-efficiency Undecanal-based N Termini EnRichment (HUNTER)^38^ using roots of 7-day-old seedlings. While the N-terminome enrichment experiment did not reveal any *bona fide* AtMC3 substrates -*i.e.* proteins cleaved after an arginine and/or lysine-, analysis of the digested proteome before enrichment provided protein abundance measurements in the different AtMC3 backgrounds (Supplementary Dataset S1). Pairwise comparisons between the different genotypes revealed a core set of proteins with their levels consistently altered (more than 50% change in abundance, t-test p-value <0.05) in the *AtMC3* overexpressing line when compared to both WT and *atmc3* mutant plants (Fig. 3A, B, Supplementary Dataset S2-3). Most of the significantly accumulating proteins in the overexpressing line was related to responses to stressful environmental conditions. In particular, four positive regulators of the response to hypoxia and oxidative stress (ACX1, At4g19880, FAD-OXR, DJ1A), one positive regulator of the response to drought stress (BGLU18), and a positive regulator of defense responses (KTI4) were found in higher abundance (Fig. 3C).Consistently, three negative regulators of ABA signaling (BGLU22, BFRUCT4, SASP), a negative regulator of drought responses (NAI2) and several peroxidases responding to oxidative stress (PER3, PER29 and PER45) were identified among the proteins depleted in the AtMC3 overexpressor line (Fig. 3C). The *atmc3* mutant was also compared to WT, but application of the selection criteria identified a single differentially accumulating protein; the chloroplastic serine peptidase CGEP, which was significantly more abundant in WT plants (Fig. S4, Supplementary Dataset S4). The lack of striking differences between WT and mutant plants could be explained by the small amount of AtMC3 naturally present in the plant and concentrated in the phloem tissue.

**Figure 3:**
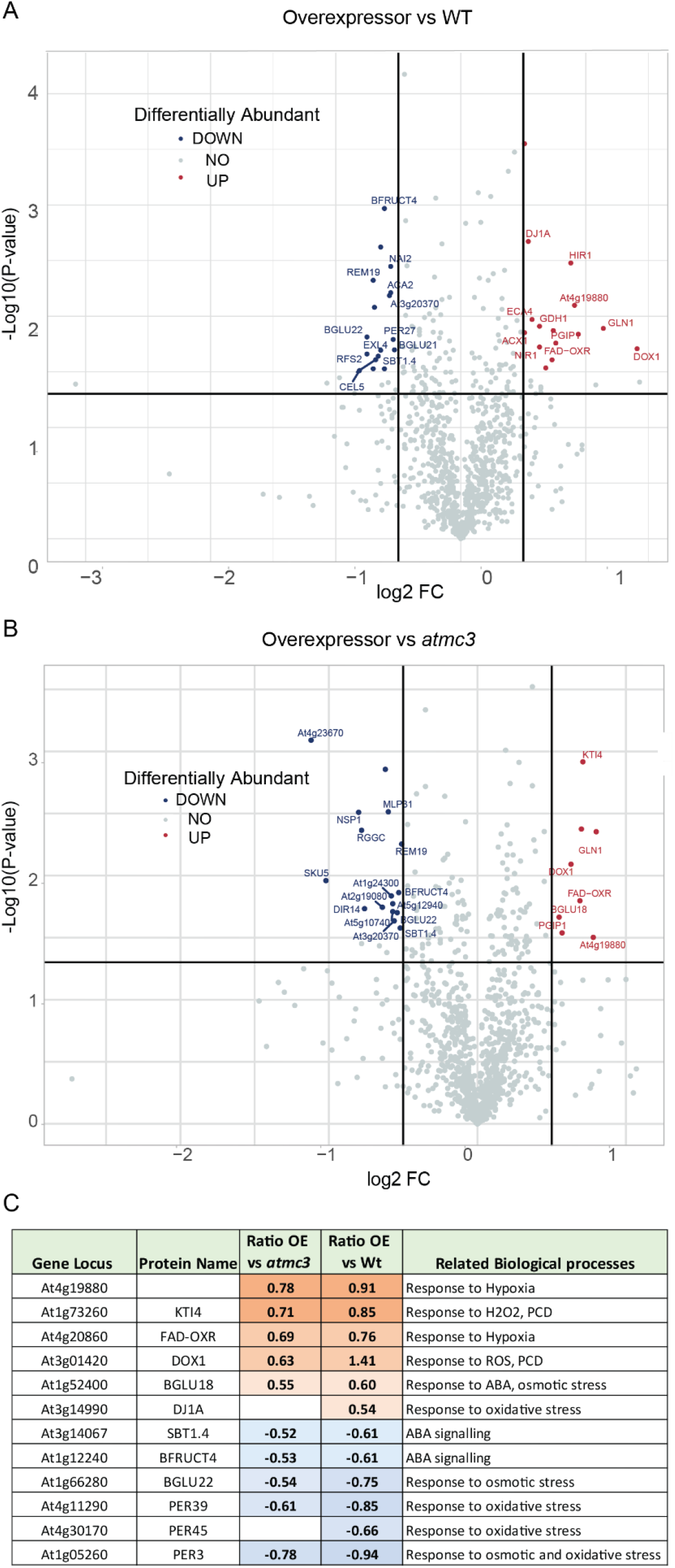
Proteomic analysis in root tissue showed differential abundance of stress related proteins depending on AtMC3 protein levels. Volcano plots of peptide abundance in (A) overexpressor line of AtMC3 compared with WT plants and (B) overexpressor line of AtMC3 compared with the atmc3 #13.3 mutant plants. N-terminal peptides with significantly reduced or increased abundance (Student’s t-test P-value<0.05 and log2 FC<–0.5 or >0.5) are highlighted with blue and red color respectively. Results are means from four independent biological replicates. (C) Summary of stress related proteins which were found differentially abundant in the comparisons performed. Biological processes are presented according to the gene ontologies (GO) following: Response to oxidative stress: [GO:0006979], [GO:0042744], [GO:1900409] Response to osmotic stress: [GO:0009269], [GO:0071472], [GO:0009651], [GO:0009414], [GO:0030104], ABA signaling: [GO:0009687], [GO:0009738], [GO:0009789], [GO:0009737], PCD: [GO:0008219], [GO:0012501], ROS: [GO:0034614], [GO:0042542]. For the complete list of proteins which were deregulated by the different expression levels of AtMC3, see Supplementary Dataset S1.

Proteomic data suggested that AtMC3 could be involved in ABA signaling and drought/hypoxia responses, both of which have a dramatic impact in the osmotic balance of the plant^39–41^. To test this hypothesis, we first examined whether *atmc3#13.3* mutant or *AtMC3* overexpressing plants displayed altered responsiveness to ABA compared to WT or to *atmc3#13.3* plants complemented with a WT copy of AtMC3 (*atmc3#13.3 pAtMC3:AtMC3-GFP*). We analyzed different processes regulated by this hormone, including germination, senescence and stomatal closure. Treatment with ABA has been shown to inhibit seed germination and seedling establishment^42, 43^. We observed that the inhibition of germination caused by different ABA concentrations was significantly less severe in *atmc3* mutants, indicating that the mutant was less sensitive to the hormone (Fig. 4A, B). In addition, seedlings lacking AtMC3 were able to establish better in media containing ABA (Fig 4B). Furthermore, since ABA is able to promote senescence in detached organs^44^, we treated rosette leaves with ABA and quantified the level of induced senescence by measuring chlorophyll levels. ABA treatment caused a less pronounced decrease in total chlorophyll in *atmc3* mutants compared to WT plants, indicating that absence of AtMC3 may affect ABA-induced senescence (Fig. 4C). Finally, since ABA is the main positive regulator of stomatal closure in response to water deficit, we tested the responsiveness of the plants with altered AtMC3 levels to ABA treatment by analyzing their ability to close their stomata in response to this hormone. As shown in Fig. 4D, stomata of *atmc3* plants were less responsive to ABA after 30 min of treatment compared to all other genotypes tested. Taken together, these results demonstrate that the absence of AtMC3 results in decreased sensitivity to ABA.

**Figure 4:**
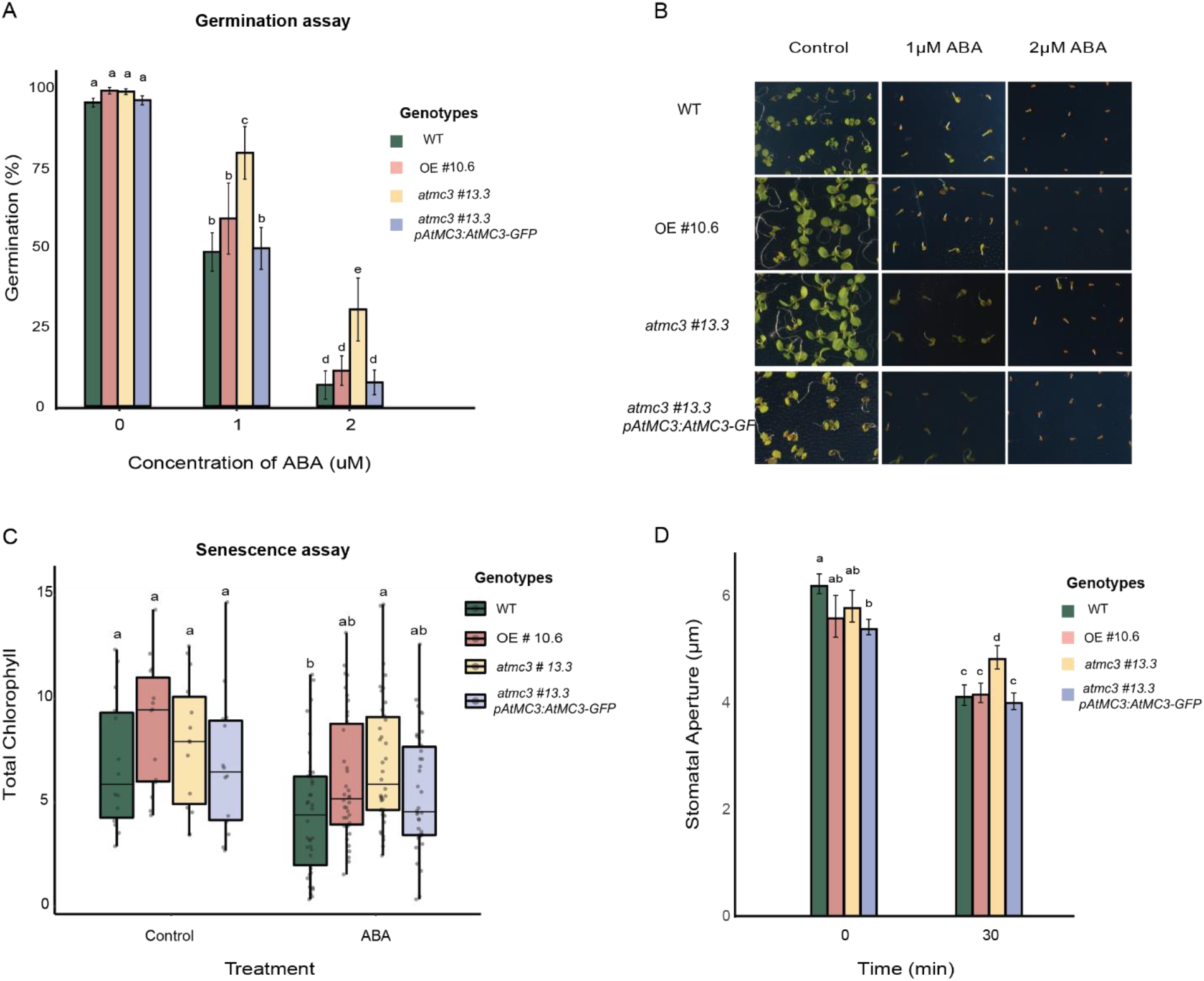
AtMC3 null mutants show less sensitivity to ABA. (A, B) Germination assays of Wt, overexpressor, atmc3 #13.3 mutant and complemented plants of AtMC3 with and without ABA treatment. Seeds were germinated on MS- medium plates supplemented with 0 (control), 1 or 2 μM ABA solution and grew for 8 days before pictures were taken (B) and germination rate was calculated (A). Bar plots in (A) are means (±SE) for five biological replicates (n> 90 per replicate/per genotype). Different letters depict significant differences in a one-way ANOVA plus a Tukey’s HSD test. (C) Senescence assay. Rosette leaves from 5-week-old plants were cut and incubated with 0 (control) and 50uM ABA. After 3 days, total amount of chlorophyll was estimated. Experiments were repeated three times and independent leaves were used to measure the absorbance (n>5 in each replicate). Different letters depict significant differences in a one-way ANOVA plus a Tukey’s HSD test. (D) Changes in stomatal aperture in WT, overexpressor, atmc3 #13.3 mutant and complementation line after ABA treatment. Cotyledons of 10-day-old plants were treated without (control) or with 10 μM ABA. Measurements of stomatal aperture in normal conditions and upon ABA treatment are given in the graph. Data are means (±SD) for three biological replicates; for each genotype, 30 guard cells were examined in each of the three replicates. Significant differences from WT were determined by Student’s t-test: *P<0.05.

### Overexpression of AtMC3 confers enhanced drought tolerance

Next, we investigated whether AtMC3 could play a role in the response to water-deficit conditions that are mainly regulated by ABA^45, 46^. For this, we first compared the survivability of lines with altered AtMC3 levels (*atmc3* mutants and *AtMC3* overexpressors) to wild type and AtMC3 complemented mutant plants to severe drought stress. Water was withheld for 8-9 days in fully-grown plants and the survival rate was calculated after re-watering. Interestingly, plants overexpressing AtMC3 (independent lines #10.6 and #5.2 tested) showed increased drought tolerance compared to WT plants (Fig. 5A, B). Furthermore, both *atmc3* mutants showed reduced survival rate under drought conditions. The predicted catalytic site did not seem to be required for the drought tolerant role of AtMC3, since a transgenic line expressing a version of AtMC3 with a cysteine to alanine substitution in the putative catalytic site in an *atmc3* #13.3 knock-out mutant background (*atmc3 #13.3 pAtMC3:AtMC3-C230A-GFP*) still restored the survival rate in a WT comparable level under drought conditions (Fig.5A, B).

**Figure 5:**
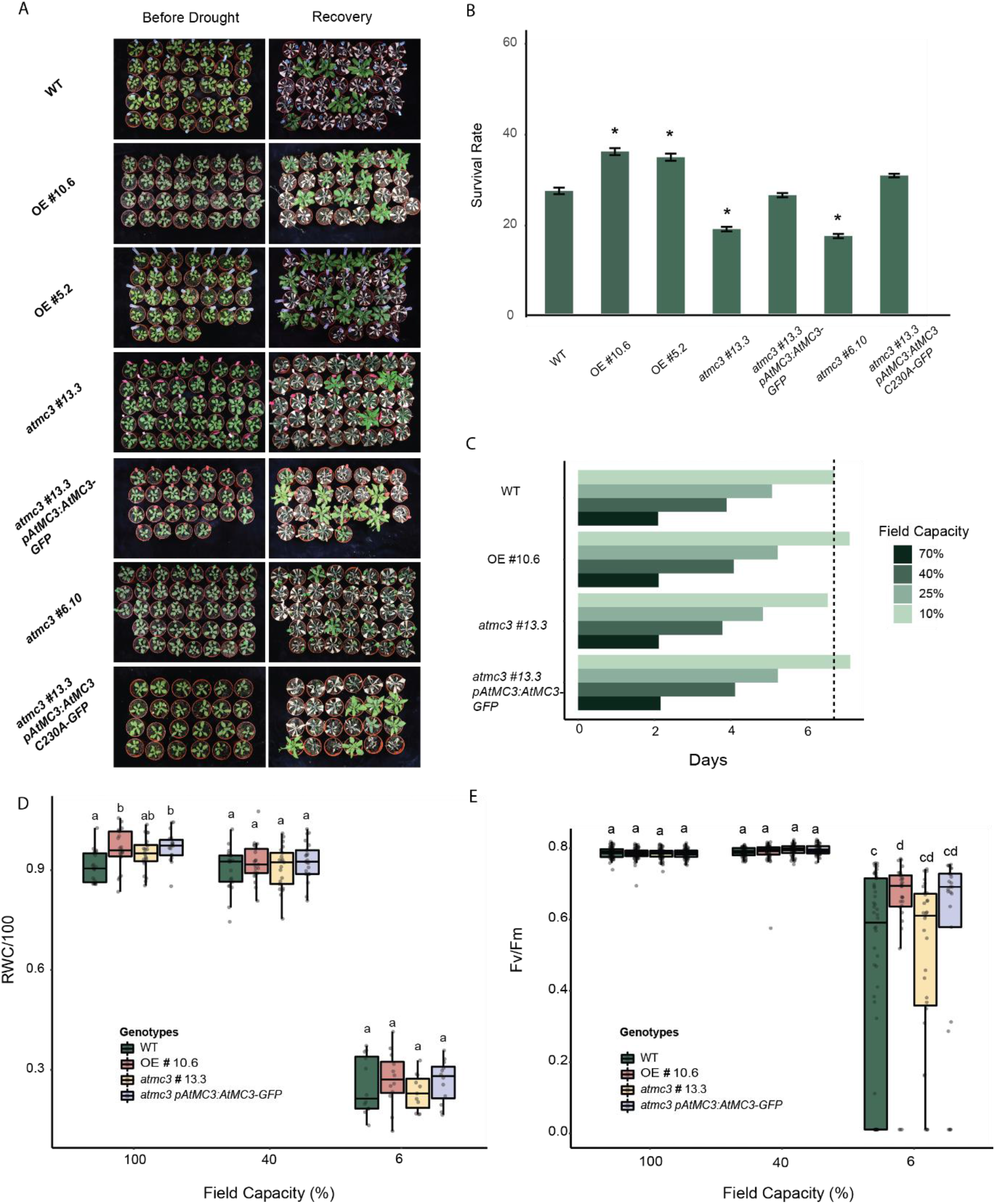
AtMC3 is involved in drought tolerance. (A) Phenotypes from 3-week-old plants before drought stress in well-watered conditions (left column) and after 5 days of re-watering and recovery (right column). From top to bottom are demonstrated 3-week-old plant rosettes of WT, OE #10.6, OE #5.2, atmc3 #13.3, complementation line of the atmc3 #13.3 mutant, atmc3 #6.10 and complementation line of atmc3 #13.3 with a predicted catalytic inactive version of the protein. (B) Plant survival rates as calculated from plants that survived after 5 days of re-watering in comparison to the total number of plants used. Averages of eleven independent biological replicates (±SE) (n >150). Asterisks indicate significant differences (p-value < 0.05) in a two-sided chi-squared test for survival ratios compared to WT. (C) Results for the days needed to reach different percentages of field capacity for WT, overexpressor, atmc3 null mutant and complementation line are shown in bar plots for five independent replicates (n >40). (D) Relative water content (RWC) of mature rosettes at 100%, 40% and 6% field capacity. Experiments were repeated four times (n >16). (E) Maximum Quantum Yield of photosynthesis (Fv/Fm) ratio for WT, overexpressor, atmc3 null mutant and complementation line at 100%, 40% and 6% field capacity Experiments were repeated four times (n>16). In (D) and (E) different letters depict significant differences within each genotype in a one-way ANOVA plus a Tukey’s HSD test.

To further characterize this response, we measured water loss through relative water content (RWC), and photosynthetic performance at specific levels of field capacity in plants subjected to drought stress, to ensure that the plants are sensing the same water loss and thus experiencing comparable stress levels. The *AtMC3* overexpressor plants along with the *atmc3* complemented line took 8-9 days to reach 10% field capacity (low water availability), whereas WT and null mutant plants needed only 7 days (Fig. 5C). This suggests that plants overexpressing *AtMC3* are able to withhold water in a more efficient way and minimize the losses while the stress is progressing, a fact that explains the higher survival detected.

Under drought progression, (40% and 6% field capacity) no differences in RWC were observed among the different genotypes. However, in well-watered conditions (100% field capacity) the overexpressor and the *atmc3* complemented line displayed slightly higher water content than the WT plants, which may indicate that they can take better advantage of environmental water when available (Fig. 5D). Maximum quantum yield of photosystem II –revealed by the ratio of variable to maximum chlorophyll fluorescence after dark-adaptation or Fv/Fm– was similar for all lines under well-watered conditions. However, when plants were experiencing severe drought, *AtMC3* overexpressing lines showed enhanced photosynthesis performance as observed from the less distributed values than WT or *atmc3#13.3* mutant plants (Fig. 5E, S5). In addition, hormonal analysis revealed that after 5 days of water deprivation, the overexpressor line contained lower levels of ABA and the cytokinin isopentenyl adenosine (IPA), which is an additional indication of reduced stress levels (Fig. S6). On the contrary *atmc3#13.3* contained elevated levels of salicylic acid (SA) and gibberellin (GA) (Fig. S6). GA levels have been shown to be reduced under drought in order to inhibit growth, aiding plants to cope better with the stress^47^. In sum, these data clearly show that increasing the levels of AtMC3 enhances the ability of the plant to face drought stress without affecting its growth rates.

The reduced ABA sensitivity displayed by the *atmc3#13.3* mutant was not caused by altered expression of ABA biosynthesis genes (*AAO3*), ABA receptors (*PYL5, PYL4*) and negative regulators of ABA signalling (*ABI1, HAB1*), as demonstrated by gene expression analysis (Fig. 6). However, the transcripts of *RD29A* and *RD22*, two dehydration responsive genes induced by ABA to maintain the osmotic balance in the cell, were downregulated in *atmc3* when no stress was applied. In addition, high levels of *ADH*, a gene known to confer tolerance to multiple stresses among these hypoxia and drought^48–50^, were detected in the *AtMC3* overexpressor under mock conditions (Fig. 6).

**Figure 6:**
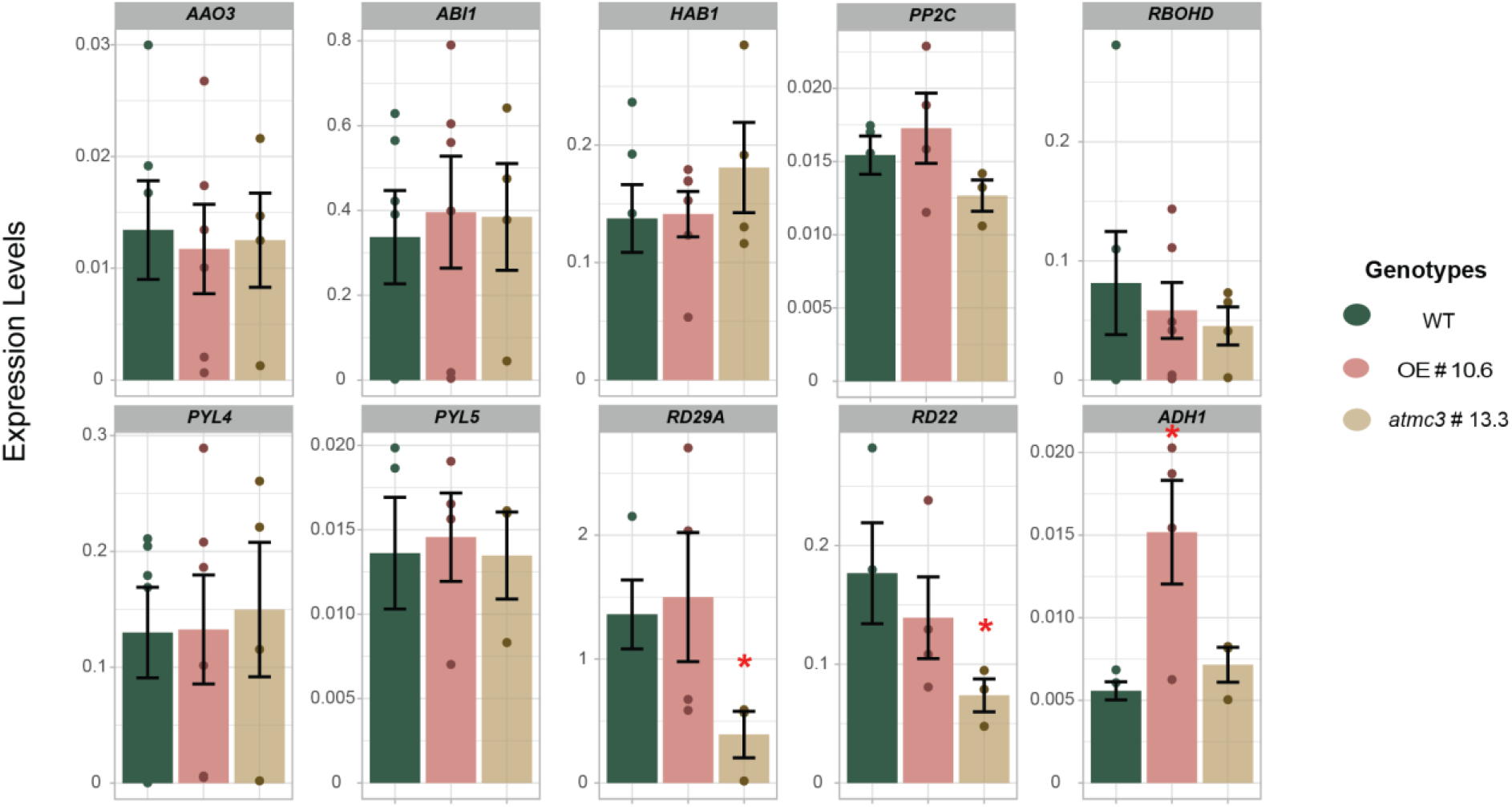
Expression levels of ABA-related genes in AtMC3 transgenic plants. The relative expression levels to the housekeeping gene EIF4a of genes related to ABA biosynthesis (AAO3), downstream signalling (HAB1, ABI1, RBOHD, PYL4, PYL5, PP2C, RD22, RD29A) and a target gene (ADH1) were analyzed from the cotyledons of 7-day-old seedlings grown in MS- media of WT, AtMC3 overexpressor and atmc3 #13.3 mutant plants. Total RNA was extracted and used for cDNA synthesis, which were then used for quantitative PCR analysis. Data are means (±SE) of three biological replicates. Significant differences from WT were determined by Student’s t-test: *P<0.05.

In an attempt to identify potential interactors of the protein, we obtained a large number of proteins that co-immunoprecipitate with AtMC3 both under mock and drought conditions (Supplementary Dataset S5). From GO analysis, in drought conditions 111 out of 600 proteins were related with metabolic processes (GO:0006091/ GO:0044281) and among those, proteins related to metabolism or transport of osmoprotectants such as sucrose and proline were enriched already in control conditions (Fig. S7A-B, Supplementary Dataset S7). In addition, many enriched proteins were related to abiotic stress-related responses (Fig. S7C), with some of them reported to show specific vascular expression such as Annexin1 (ANN1) and ABCG11 transporter. ANN1 is responsible for the post-phloem distribution of sugar in the root tip and when overexpressed, it confers drought tolerance^51, 52^. Aquaporins, dehydrins and oxidative stress-related proteins were enriched under both control and drought conditions (Fig. S7C). Finally, we detected cell wall modification enzymes such as hydrolases and cellulose synthases that immunoprecipitated with AtMC3 exclusively under drought conditions (Supplementary Dataset S6).

To determine whether AtMC3 is involved in responses to other osmotic-related stresses, we subjected *atmc3#13.3* mutant, AtMC3 overexpressor and WT plants to hypoxia and salinity conditions. Under low oxygen conditions (hypoxia) the AtMC3 overexpressing line showed a markedly higher survival rate compared with WT plants and similar to the *prt6* mutant that constitutively induces hypoxia responsiveness through the PCO N-degron pathway^53^ (Fig. S8A). Interestingly, the *atmc3* #13.3 mutant also showed enhanced survival rates under hypoxia conditions. Several hypoxia-marker genes were tested under basal conditions. A clear upregulation of *ADH1* under mock conditions in the AtMC3 overexpressing line compared to WT could be an indication of a better/faster response to hypoxia (Fig. S8B). On the other hand, increased levels of salinity result in inhibition of primary root growth and induce lateral root formation. Plants overexpressing AtMC3 produced a larger number of lateral roots than WT in response to salinity (Fig. S8C), a phenotype usually observed in plants that display higher salinity tolerance. In contrast, increased levels of salinity resulted in similar levels of primary root growth inhibition in all lines tested (Fig.S8D). In conclusion, our data indicate that AtMC3 may play a role in the responses of the plant to stresses that cause osmotic imbalance to the plant resulting from either excess or absence of water.

### Increased AtMC3 levels lead to early metaphloem development and maintain functional transportation under osmotic stress

Considering the specific spatial expression and localization pattern of AtMC3 and its involvement in responses to osmotic stress conditions, we next investigated whether stress impinges on the developmental formation of phloem tissues and whether AtMC3 plays a role in these conditions. Upon osmotic stress, root cells lose water content and accumulate sugars and other osmoprotectant compounds to balance the water shortage from the environment, minimize losses and avoid cellular damage^15, 54^. As previously documented, incubation of WT plants in media containing sorbitol to cause osmotic imbalance resulted in root growth arrest and a comparable decreased meristematic activity^55^, a phenotype also observed in all genotypes tested (Fig. 7A).

**Figure 7:**
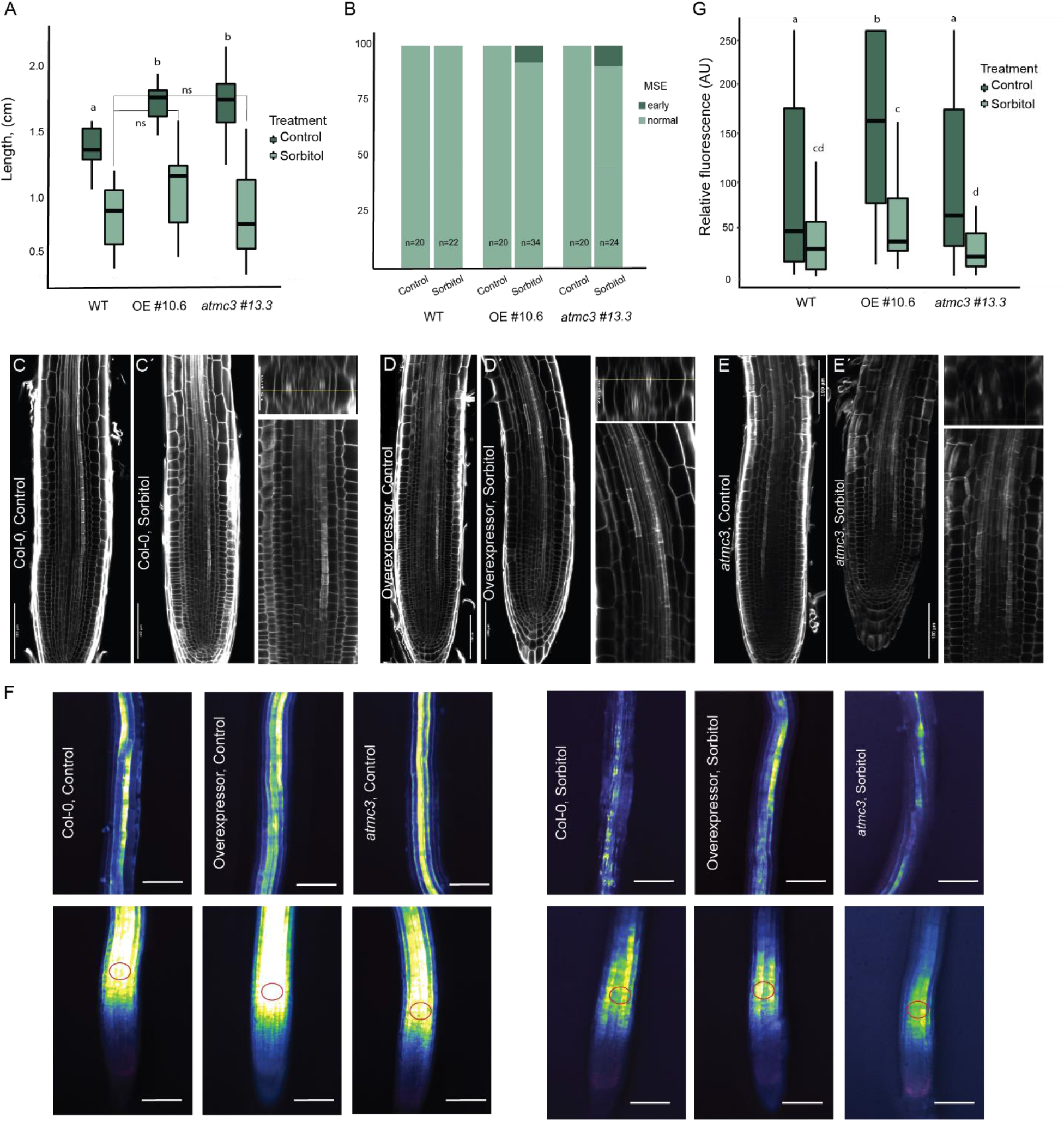
Vascular tissue formation and function under osmotic stress is affected by AtMC3 levels. (A) Root length of 6-day-old seedlings measured from WT, overexpressor, atmc3#13.3mutant plants in control (darker green) and under 120 mM sorbitol treatment (lighter green). (B) Quantification of the early (darker green) metaphloem sieve elements (MSE) that appeared differentiated earlier that protoxylem strands and displayed in (C-É). (Ć, D′, É) For sorbitol treatments, on the right lower panel is the magnification of the vascular tissue presented on the left panel. Transverse observations of the root are also present on the right upper panel. Calcofluor staining was used to visualize the roots. Scale bar 100μm. (F) Representative pictures from 6-day-old seedlings for WT, overexpressor of AtMC3 and atmc3 #13.3 mutant in control and sorbitol conditions. Upper panels are showing the successful loading of the phloem tissue and in lower panels are presented the root tips approximately 1 h after loading CFDA on cotyledons. CFDA fluorescence is visualized in LUT Green Fire Blue; fluorescence intensity was quantified in the region indicated by red circle. G) Fluorescent intensity quantification for the phloem unloading zone in WT, overexpressor AtMC3 and atmc3 #13.3 mutant in 6-day-old seedlings of control and sorbitol treated plants; Scale bar 250μm; letters depict significant differences in a one-way ANOVA plus a Tukey’s HSD test.

To better assess whether sorbitol affects phloem development, sorbitol-treated roots were stained with calcofluor and analyzed by confocal microscopy. A prolonged exposition (72 h) to osmotic stress triggered a premature differentiation of PSEs, as indicated by the appearance of stained cell walls close to the quiescence center (Fig. 7C-E′). While this phenotype is common to all tested genotypes, 10-20% of the roots with altered AtMC3 levels showed a premature differentiation of MSE as manifested by their thick cell wall, even before the differentiation of protoxylem strands (Fig 7B, C-É). Metaphloem is the main transport form of the phloem, which functionally replaces the protophloem tissue when the surrounding cell types have already differentiated. Although the two phloem tissues share the same precursor cell, metaphloem formation usually occurs in the maturation zone, approximately at 27 nm above the QC^16, 56^. These data suggest that osmotic stress may result in earlier phloem differentiation to preserve the osmotic balance and AtMC3 levels specifically affect metaphloem differentiation. To elucidate how long distance transportation is affected under osmotic stress conditions, we used 5(6)-carboxyfluorescein diacetate (CFDA), a known phloem-mobile probe^57^. Seedlings were treated with sorbitol for 72 h and CFDA was then applied for 1 h on the cotyledons to ensure its successful loading into the vasculature. Interestingly, we observed that the overexpressing AtMC3 line displayed higher amounts of CFDA in the protophloem unloading zone^57^ compared to the rest of the genotypes even under control conditions. Under osmotic stress, *AtMC3* overexpressors were able to maintain significantly higher levels of transport compared to the *atmc3* mutant plants (Fig. 7F, G). To sum up, fine-tuned levels of *AtMC3* are required for the plants to show the appropriate sensitivity to drought stress by enhancing vascular formation; increased levels of the protein to ensure proper function of the tissue.

## Discussion

The Arabidopsis type I metacaspase *AtMC3*, exhibits a vascular-specific expression pattern concentrated in the phloem pole (Fig 1 and ^19^). Our data show that *AtMC3* displays a restricted expression pattern with a shift from the differentiating PSE to the surrounding CCs (Fig. 1, S2), similar to multiple genes related to phloem formation such as *APL*, *NAC45/86* and *CLE45*^22, 58^. Further, we demonstrate that the AtMC3 protein also localizes in the vasculature.

Although altered levels of the protein did not cause any detrimental changes in plant growth and development (Fig 2, S3), proteomic analysis suggested that AtMC3 is involved in abiotic stress responses. ABA downstream signaling, osmotic and hypoxia stress-related proteins appeared deregulated in plants with altered AtMC3 levels, even under control growth conditions (Fig. 3). Multiple peroxidases accumulated to lower levels in plants overexpressing *AtMC3*, potentially indicating changes in the endodermal layer of the root. In contrast, other proteins related to protection against oxidative stress, such as ACX1 or DOX1 overaccumulated in the overexpression line. In addition, BGLU22, which shows reduced expression upon mannitol treatment^59^, accumulated to higher levels in the mutant, while BGLU18 (or ATBG1), involved in the *de novo* biosynthesis of ABA^60^, showed the opposite behavior. Supporting this data, BGLU18 was found also to co-immunoprecipitate with AtMC3 under drought stress (Fig. S7C). Interestingly, SASP, a protease that negatively regulates drought responses by leading OST1 to degradation^61^, was found less abundant in the *AtMC3* overexpressing line. Similarly, we detected reduced accumulation of NAI2, which is involved in ER body formation and was shown to interact with the membrane-associated protein At14a-like1 (AFL1) under stress conditions enhancing downstream signaling^62^. Reduction of NAI2 could be an indication of increased proline accumulation, which assists as osmoprotectant and improves responses to drought.

ABA has been long considered a “stress” hormone, as it regulates most of the plant responses to challenging environmental conditions. Plants lacking AtMC3 showed reduced sensitivity to ABA treatment (Fig. 4), which explains their diminished ability to cope with drought stress and the fact that they lose water faster than WT plants (Fig. 5B, C). Stomata respond to elevated ABA levels by closing immediately to avoid excess loss of water when conditions are not favorable. Plants null for AtMC3 displayed a delay in their responsiveness to the hormone, which could have detrimental consequences for their survival under natural environmental conditions (Fig. 4C). Reduced expression of specific dehydration-responsive genes in *atmc3* mutants could indicate that AtMC3 may be involved in signaling downstream ABA (Fig. 6). Alternatively, the limited responsiveness to ABA observed in the *atmc3* mutant could also suggest that AtMC3 directly or indirectly participates in long-distance transport of the hormone, mediated by the xylem.

We demonstrated that plants overexpressing AtMC3 were more tolerant to drought stress, as indicated by their increased survival rate and their ability to maintain their photosynthetic capacity more stable in high levels of water loss (Fig. 5, Fig. S5). In contrast, *atmc3* mutants showed reduced survival compared to WT plants, which was restored by complementing the mutant both with a WT copy of AtMC3 or with a variant version carrying a mutation in the predicted catalytic site. The fact that the predicted protease function of AtMC3 is not required for its function raises the question of whether the protein might act as a potential stress sensor in CCs. This notion is underscored by the fact that we were not able to detect any direct AtMC3 proteolytic cleavage from the degradome analyses. In fact, metacaspases have been proposed to act as homeostatic rheostats, in some cases with functions partly independent of their catalytic activity^63^.

The vascular tissue is of great importance for the establishment of inter-organ communication throughout the plant body. The interaction between phloem and xylem and the maintenance of tissue plasticity is key for the plant adaptation to stress. Sugars are actively loaded into the phloem from source tissues and the xylem is transferring water into the phloem strands to generate the appropriate pressure and flow of the macromolecules though the vasculature. Upon arrival to the sink tissues, sugars will be unloaded and distributed and water returns into the xylem files^15^. Osmotic stress affects the source-sink relationships, and the phloem needs to adjust the osmotic pressure generated from water reduction in the xylem files. An efficient way to achieve this could be increasing the levels of osmoregulators in the phloem sap^15^. However, little is known about signal perception and the responses of phloem tissue under drought stress conditions^64^. Genes involved in vascular development were shown to be affected under osmotic stress^65^, which led to the speculation that promoting phloem development could increase tolerance. Indeed, we showed that when faced with osmotic stress, plants form protophloem strands closer to the quiescent center (Fig. 7C-E′). Additionally, altered AtMC3 levels result in premature differentiation of metaphloem cells (Fig. 7B). A functional metaphloem closer to the root tip as observed in the *AtMC3* overexpressing line could provide an advantage to the plants by facilitating the allocation of photoassimilates and osmoprotectant molecules to decrease the water potential in roots or provide the necessary energy for the surrounding tissues to maintain their function and respond. This fact was further supported by the enhanced long-distance transport of CFDA to the root tips in *AtMC3* overexpressor plants (Fig 7G). Osmotic balance may be maintained more effectively along with the simultaneous induction of xylem differentiation by ABA^14^. Counterintuitively, the premature metaphloem differentiation observed in the *atmc3* mutant, did not result in a similar increase in conductivity, which may suggest that the correct dosage of AtMC3 is required to fine-tune the trade-off between osmotic protection and root development. Faster protoxylem development could lead to increased lethality in early seedling stages due to osmotic stress, caused by reduced cell wall viscoelasticity, as was shown recently^66^. In contrast, additional xylem strands generated in later stages can provide tolerance^14^. Specifically, under drought stress, proteins related to cell wall modification and phloem development (such as CALS8 and SEOB) co-precipitated with AtMC3, a fact that highlights the importance of the plasticity of the cell borders for the plant responses (Supplementary Dataset S6).

Interestingly, we observed that AtMC3 may also participate in hypoxia and salinity responses, which result in osmotic imbalances from excess or absence of water, respectively (Fig S8A, S9). Higher *ADH1* expression levels observed in *AtMC3* overexpressors at the seedling stage (Fig.6, S8B) may give these plants an advantage to cope with multiple stresses^48, 50^ similar to the *prt6* mutant that constitutively enhances *ADH1* and other hypoxia-related gene expression^53^.

The specific phloem localization of AtMC3 raises the question of whether this protein might be acting as a sensor of stress conditions and what is the role of CCs in signal perception. The phloem can transport signaling molecules when plants are under stress, as it is the case with the CLE25 peptide, which is translocated from the root to the shoots under drought and induces ABA synthesis and stomatal closure^67^. In the case of AtMC3, the supporting cell types of proto- and metaphloem could be the orchestrators for the decisions of cell differentiation – leading to enhanced proto- and metaphloem development and energy allocation. Hormones are also involved in the complexity of drought responses. Overexpression of the vascular specific-localized brassinosteroid (BR) receptor BRL3 was shown to confer drought tolerance without affecting plant growth, driving the accumulation of osmoprotectant molecules to the root^68^. In this context, multiple GRF proteins (GRF1,2,10) as well as BSK1, which belong to BR downstream signalling pathway^69^ co-immunoprecipitated with AtMC3. It would be of interest to investigate the relationship between hormones and AtMC3 in order to elucidate the tissue-specific regulation of downstream responses.

Overall, our results document the role of a vascular metacaspase in osmotic stress responses through a yet to be defined mechanism. It is important to highlight the fact that AtMC3-overproducing plants are able to maintain WT-like growth and yield, unlike other drought resistant plants reported to date. Food sustainability and agriculture are threatened by incremental climate instability and water scarcity. Severe drought events result in significant reduction of dry weight and lead to diminished harvested biomass and yield. Therefore, the elucidation of genetic features that may allow tissue specific engineering to enhance drought tolerance without taking a toll from plant growth may constitute an extremely valuable strategy to sustainably improve valuable crops.

## Methods

### Plant material and growth conditions

All experiments were performed using *Arabidopsis thaliana* (L.) Heynh. accession Col-0. Seeds were surface-sterilized with 35% NaClO. and shown in agar Murashige and Skoog (MS) media with vitamins and without sucrose. After 48 h of vernalization, plates were grown vertically under long day (LD) conditions (16 h light/8 h dark; 22 °C, 60% relative humidity) unless indicated otherwise. Genotypes used in this study are described in Supplementary Table 1. DNA C-TAB extraction protocol^70^ was used for all plant genotyping experiments. Primers used for genotyping can be found in Supplementary Table 2.

### DNA constructs

To obtain the reporter line *pAtMC3:GUS-GFP* Gateway cloning was used. Briefly, a 1500 bp promoter region of *AtMC3* was amplified with primers pAtMC3_F/R and cloned into the binary vector pBGWFS7^71^. *UBQ10:AtMC3-GFP, pAtMC3:AtMC3-GFP, UBQ10:AtMC3-C230A-GFP* and *pAtMC3:AtMC3-C230A-GFP* constructs were generated using the GREENGATE cloning strategy as described previously^72^. Briefly, *AtMC3* promoter fragment (∼2500bp) was inserted in the pGGA entry module, whereas UBQ10 in pGGA was commercially available from Addgene. The full-length genomic DNA sequence of *AtMC3* was cloned into the entry vector *pGGC*. The CDS sequence of AtMC3 was inserted in *pGGB* module. To generate the catalytic inactive version of the gene, the Quick site mutagenesis kit (Agilent Technology) was used with *pGGC*/*B_MC3* module as a template. For the overexpression lines *pGGA-UBQ10,* p*GGB*- *dummy, pGGC-AtMC3* genomic or *pGGC- AtMC3 C230A* genomic, *pGGD-eGFP*, *pGGE-UBQ10terminator*, *pGGF-BASTA* were combined in the *pGGZ003*-empty destination vector. For the expression of the protein under the native promoter *pGGA*-promoter *AtMC3*, *pGGB-AtMC3* CDS sequence, *pGGC-eGFP*, *pN95 (RBCSt.D-F-pGGD000*), *pGGF Alli-YFP* (green fluorescent seed plant selection) combined in the *pGGZ003*-empty destination vector. Plants were transformed using *Agrobacterium tumefaciens* (GV3101)-mediated floral dip as previously described^73^. Homozygous transgenic lines were selected either on MS media supplemented with 20 μg ml^−1^ Basta (glufosinate-ammonium) or by red/green fluorescence depending on the selection included in the destination vector used. Primers used for cloning can be found in Supplementary Table 2.

### CRISPR mutagenesis

For generation of the *atmc3* CRISPR mutants the target sequence was selected using CRISPOR program^74^. DNA backbone fragments of a total length of 500 bp contained: 20 bp of the target sequence neighboring the PAM (3 bp Protospacer Adjacent Motif) sequence, overhang *attB* sequences for Gateway cloning, *tracrRNA* sequence, *U6* promoter sequence and restriction sites for the cloning were ordered as gBlocks® from IDT. Three μl (50ng/ul) of the *gBlock* sequence, was introduced to 1µl (150ng/µl) of *pDONR207* with BP reaction using 1μl of BP clonase II enzyme (Thermo Fisher Scientific). For the combination of three gRNAs in one vector, the vectors including the individual gRNAs were digested with restriction enzymes BamHI/PstI/SalI. The *pDONR207* vector containing the triple gRNAs sequences and the single gRNA sequence was then transferred into the *pDe-CAS9-DsRED* vector^75^ via LR reaction (Thermo) following the manufacturer’s protocol. The destination vector was delivered to WT plants using *A. tumefaciens* (GV3101)-mediated floral dip as previously described^73^.

### Confocal analysis

Confocal laser scanning microscopy was performed using a Leica SP2 AOBS inverted confocal microscope (Leica Microsystems, Wetzlar, Germany) or a Zeiss LSM 780. eGFP fusion was excited with a 488 nm Argon laser and detected using a 505–530 band pass emission filter. Propidium iodide staining was excited using a 561 nm He-Ne laser and detected using a custom 595–620 nm band pass emission filter. Calcofluor stained samples^76^ were excited at 405 nm and emission was collected at 425-475 nm. Different Z stacks and transverse optical versions were obtained using Image J software.

### Cotyledons vein pattern

Cotyledons were removed from 6-day-old seedlings and submerged for 1 h in a 3:1 95% ethanol: acetic acid solution at room temperature. Subsequently, they were washed twice for 30 min with 70% ethanol and were incubated overnight at 4°C with 100% ethanol. 10% NaOH was added for 1 h and the samples were left at 37°C before Clear-See was added as a final step. Digital pictures were obtained on a Leica DM6 epifluorescence microscope (Leica, Wetzlar, Germany).

### GUS (β-glucuronidase) staining

Seedlings were harvested from plates at the 2-cotyledon stage, then fixed/stained according to iced-acetone fixation method. Incubation with GUS staining solution containing 1 mM X-Gluc, 2.5 Mm K_3_Fe(CN)_6_, 2.5 mM K_4_Fe(CN)_6_ and 0.1% Triton x-100 in 100 mM sodium phosphate buffer (Na_2_HPO_4_/NaH_2_PO_4_, pH 7.0), was overnight at 37°C, followed by EtOH clearing for 5 minutes Five seedlings were mounted in 50% glycerol and were viewed at 10X at the root-tip, hypocotyl, shoot apical meristem, and cotyledons. Mature plants were harvested from soil. Hypocotyls were stained with GUS solution, fixed in FAA (50% ethanol, 10% formaldehyde, 5% acetic acid), embedded in LRwhite+PEG and sectioned at 18 µm with a microtome after embedding in Entellan.

### Transcriptional Analysis

For quantitative PCR expression analysis, tissue was collected separately from roots and cotyledons of 7-day-old seedlings. RNA was extracted with the Maxwell RCS Plant Kit (Promega) according to the manufacturer’s instructions followed by DNAse I treatment. cDNA was synthesized from 2 µg of RNA template using the High Capacity cDNA Reverse Transcriptase Kit (Applied Biosystems). For the qPCR reaction mixture KAPA SYBR FAST Master Mix (2x) Universal (Kapa Biosystems) was used with a final concentration of 10 ng template cDNA. Measurements of CT values were obtained on a LightCycler 480 (Roche Applied Science). Primer performance was checked by efficiency and melting curve (Applied Biosystems 2004). *EIF4a* (AT4G18040) was used as the reference housekeeping gene and calculations of the relative transcription level were done with the comparative 2−CT -method^77^. All primers are found in Supplementary Table 3.

### Proteome and Terminome analysis

Four independent pools of WT, *atmc3#13.3* mutant and *AtMC3* overexpressor line seedlings were grown on MS- media plates for 7 days. Approximately 600 mg of roots were separately harvested, ground in a fine powder with liquid nitrogen and resuspended with 1 ml of Guanidine hydrochloride extraction buffer (6M GuaHCl, HEPES 1 M, EDTA: 5 Mm, adjusted to pH 7.5) supplemented with protease inhibitor cocktail (Roche). Homogenates were filtered through a 100 µm cotton mesh, centrifuged at 500 *× g* for 5 min at 4°C, and the filtrate was centrifuged at 10 000 *× g* for 1 min at 4°C. Proteins in the supernatant were precipitated with chloroform/methanol and resolubilized in 6 M GuHCl, 100 mM HEPES pH 7.5. Protein concentration was estimated using the BCA assay at 2μg/μl (Bio-Rad) and frozen and −80°C until further processing. HUNTER N-terminome analysis was performed as previously described^38^.

### Total Protein Isolation and Immunoprecipitation

WT and *AtMC3* overexpressor plants were grown for 3 weeks before subjected to water withdrawal. Tissue was collected from well-watered plants (control condition) and plants subjected to drought for 5 days and frozen immediately with liquid nitrogen. Protein extraction buffer (20 mM Tris-HCl, pH 7.5, 150 mM NaCl, 1mM EDTA, Glycerol, 1% Triton X-100, o,1% SDS and 5mM DTT) supplemented with a 1:100 dilution of plant protease inhibitor cocktail (Roche) was added to the ground samples and centrifuged 20 minutes at 6,000 *x g* at 4°C. The supernatant was collected and centrifuged at 11,000 *x g* 4°C. The supernatant was again collected and protein concentrations were measured using Bradford (BIORAD) and equalized at 1.5μg. Fifty microliters of anti-GFP magnetic beads (Miltenyi Biotec) were added to the protein extracts and incubated for 1 h at 4°C in a rotary mixer. Magnetic beads were immobilized on a magnetic separator (Miltenyi Biotec) and washed and eluted according to the manufacturer’s instructions. Three independent plants were bulked in each of the four independent biological replicates (*AtMC3* overexpressor and WT plants).

### LC/MS/MS and Data Analysis

LC-MS/MS analysis was performed with an UltiMate 3000 RSCL nano-HPLC system (Thermo) online coupled to an Impact II Q-TOF mass spectrometer (Bruker) via a CaptiveSpray ion source boosted with an ACN-saturated nitrogen gas stream. For preHUNTER samples, peptides were loaded on an Acclaim PepMap100 C18 trap column (3 µm, 100 Å, 75 µm i.d.×2 cm, Thermo) and separated on an Acclaim PepMap RSLC C18 column (2 µm, 100 Å, 75 µm i.d.×50 cm, Thermo) with a 2 h elution protocol that included an 80 min separation gradient from 5% to 35% solvent B (solvent A: H2O+0.1% FA, solvent B: ACN+0.1% FA) at a flow rate of 300 nl min–1 at 60 °C. For IP samples, peptides were loaded on a µPAC pillar array trap column (1 cm length, PharmaFluidics) and separated on a µPAC pillar array analytical column (50 cm flowpath, PharmaFluidics). Line-mode MS spectra were acquired in mass range 200–1400 m/z with a Top14 method.

For analysis of preHUNTER and IP data, peptides were identified by matching spectra against the UniProt *Arabidopsis thaliana* protein database (release 2020_2) with appended MaxQuant contaminant entries using the Andromeda search engine integrated into the MaxQuant software package (version 1.6.10.43) with standard settings^78^. Further analysis was performed using the Perseus software package (v 1.6.14.0 or v 1.6.15.0). Only proteins quantified in at least two of the four biological replicates were used for pairwise comparisons of each of the conditions. Protein ratios were median-normalized within each replicate before assessing differential expression with a one-sample t-test as implemented in Perseus. For preHUNTER proteins changing at least 50% in abundance (log2 fold change < –0.58 or >0.58) supported by a t-test p-value <0.05 were considered as differentially accumulating. For AtMC3 IP analysis proteins flagged as contaminant or reverse protein entries were excluded from analysis, as were proteins not quantified in at least three of the four biological replicates in at least one of the four experimental groups (WT and *AtMC3*-overexpressor plants grown in wild type and drought conditions, respectively). Missing values were imputed using Perseus standard settings, and significant differences between the individual experimental groups followed by analysis for significant changes using the ANOVA test (p-value 0.05) followed by Tukeýs Post-Hoc-test (FDR<0.05).

### ABA sensitivity assays

More than 100 seedlings were sown on MS plates with or without ABA at concentrations of 1 or 2 μM. Plates were stratified for 2 days at 4°C and then transferred to a controlled growth chamber. Germination was quantified after 8 days as the number of seeds that showed both radical protrusion and elongation.

For senescence measurements, detached cauline leaves from 5-week-old plants were soaked in distilled water without or with 50 μM ABA for 3 days. Each leaf was then weighed and mechanically homogenized. Two ml of 80% acetone were added and samples were incubated at 4°C for 16 h in dark. The concentration total chlorophyll was calculated as described before^79^.

To measure stomatal aperture, cotyledons were collected from 10–days-old seedlings and immediately immersed in a stomata-opening buffer with 10 mM KCl, 0.2 mM CaCl_2_, and 10 mM Mes-KOH (pH 6.15) for 2^1/2^ h under continuous white light at 22°C. ABA was then added into the buffer to a final concentration of 10 μM and the cotyledons were collected after 30 min. Guard cells were photographed using a LEICA DM6 microscope, and stomatal widths were measured with Image J software.

### Drought stress for scoring plant survival

Drought survival rate was measured as described^68^. Briefly, seedlings grown in MS- agar plates were transferred after 7 days to individual to pots containing 30 ± 0.5 g of substrate (plus 1:8 v/v vermiculite and 1:8 v/v perlite). For each biological replicate, 40 plants of every genotype were grown for 3 weeks before subjected to severe drought stress. Water was withheld for 8 days followed by re-watering. After the 6-day recovery period, the surviving plants were counted and photographed (statistical analysis with two-sided chi-squared test, *p*-value < 0.01). Every genotype was tested at least in five independent replicates.

### Physiological parameters and chlorophyll fluorescence

Seedlings grown in MS- agar plates for 7 days were transferred into individual pots containing 30 ± 0.5 g of substrate (plus 1:8 v/v vermiculite and 1:8 v/v perlite). Plants were grown for 3 weeks before being subjected to water withholding, while well-watered plants were used as control. Relative Water Content (RWC, %) was calculated according to the formula: (FW-DW)/(TW-DW). Fresh weight (FW) was obtained by harvesting and weighing freshly detached rosettes. Turgid weight (TW) was obtained by putting cut rosettes into a tube with de-ionized water for 16 hours at room temperature, removing excess water by wiping with absorbent paper and weighing plant material. Rosette dry weight (DW) was recorded after an overnight incubation at 80°C in a dry oven. The time course drought stress assay was started by withholding water until reaching 30%, 60%, 75%, and 90% water loss. Pots were weighted daily to monitor evapotranspiration (CW). Water saturated soil for every pot was weighted initially (SW) and several pots with soil were placed in the oven for 72h to calculate the dry weight (DW). Field capacity was calculated for each pot daily by the formula (CW-DW)/(SW-DW). Drought-related experiments were repeated four times and at least four plants per genotype and treatment were used in each experiment.

Photosynthesis efficiency was measured in control (well-watered) and drought-treated plants at 40% and 6% field capacity. After 30 min of dark adaptation, the kinetics of chlorophyll fluorescence in whole rosettes were monitored by measuring F0 in the dark and Fm with initial saturation pulse using Imaging PAM M-series, MAXI version device. Fv/Fm and Fv′/Fm′ (PSII efficiency) ratio for the maximum quantum efficiency upon dark and light conditions was calculated according to the manufacturer’s instructions (Walz, Effeltrich, Germany).

### Osmotic stress assays

For salinity stress experiments, seeds were sown on MS- medium, stratified for 2 days at 4°C and grown for 3 days. Then, they were transferred to MS- plates containing increasing NaCl concentrations (0 for control, 50 and 100 mM) for 7 additional days. Primary root length was measured 10 days after germination using Image J software. Number of emerged lateral roots was counted using an Olympus DP71 stereomicroscope. Four independent replicates were performed.

For sorbitol treatments, seeds were sown on ½ MS medium supplemented with 1% sucrose, stratified for 2 days at 4°C and grown for 3 days in continuous light conditions. Then, they were transferred to ½MS plates without sucrose containing 120 mM sorbitol for 3 additional days before confocal microscopy analysis. Primary root length was measured at 6 days after germination using Image J software.

### CFDA

A stock of 5-(and-6)-carboxyfluorescein diacetate (CFDA) was prepared by dissolving 5 mg/ml CFDA (Sigma Aldrich) in acetone. For loading, 10x CDFA dilution was prepared fresh with dH2O. Plants were grown vertically for 3 days prior to transfer in 120 mM sorbitol media for 3 additional days before the treatment. A droplet of CFDA was supplied to the cotyledons of 6-day- old seedlings after pinching them with fine tweezers. Fluorescence intensity was imaged after 1 h of incubation at identical settings for all roots using a Leica stereomicroscope with GFP filter and was calculated as mean intensity values from a circular region in the area above the meristematic zone of the root tip using Fiji/ImageJ. Experiments were performed in four independent replicates (n>30).

### Hypoxic stress assays

Hypoxia stress assays were carried out as previously^80^. Seedlings were grown in ¼ strength MS plates for 4 days under short day (Conditions: T= 20°C, Light intensity= 110 µmol m^-2^s^-2^). Plates without lids were then transferred to a desiccator (hypoxia chamber) in the dark and nitrogen gas was introduced at a flow rate of 4 l/min, for 4 h. During this time oxygen within the chamber was measured and shown to reduce to less than 0.3% within 1 h. Control samples were placed in a similar desiccator under dark conditions but with air introduced (normoxia). After 4 h plates were removed, sealed with micropore tape and growth of the root tips was marked before transferring them back to the growth room for recovery. Extension of primary root tips beyond the marked point was scored as root tip survival for each seedling after 3 days of recovery.

### Hormone quantification

Three-week old plants were subjected in drought stress by water withholding for 5-6 days. Approximately 100 mg of leaf material were harvested from plants under drought and well-watered conditions and ground to a fine powder in liquid nitrogen. IAA, ABA, SA, JA, GA1, GA3, GA4, GA7, ACC, tZ, tZR, OPDA, IPA and melatonin were quantified as described previously^81^.

### Data availability

The mass spectrometry proteomics data have been deposited to the ProteomeXchange Consortium^82^ via the PRIDE partner repository^83^. Quantitative proteome data can be accessed with the dataset identifier PXD035957, IP-MS data with the dataset identifier PXD035975.

## Acknowledgements

We kindly thank Marc Planas for starting the work and Fabien Nogué for sharing CRISPR constructs and expertise. We are thankful to Moritz Nowack for sharing resources and equipment. We also thank Miguel Ángel Moreno-Risueño and David Blasco-Escámez for inspiring discussions and sharing expertise, and Paula Muñoz and Serveis Científico-Tècnics from the University of Barcelona for technical assistance in hormone quantification. Research in the NSC- MV lab is funded by project PID2019-108595RB-I00 funded by MCIN/AEI/ 10.13039/501100011033 and and through the “Severo Ochoa Programme for Centres of Excellence in R&D” (SEV-2015-0533 and CEX2019-000902-S funded by MCIN/AEI/ 10.13039/501100011033). This work was also supported by the CERCA Programme/Generalitat de Catalunya. EP and JS-L were supported by fellowships BES-2016-077242 and BES-2017- 080210, respectively, funded by MCIN/AEI/ 10.13039/501100011033 and by “ESF Investing in your future”. NR and RdP were supported by fellowships FPU2019-03778 and FPU2018-03285, respectively, funded by Ministerio de Universidades. The AR-V lab was funded by the Swiss National Foundation, Stavros-Niarchos/ETH-Foundation (EK) and a Swiss government fellowship (RRT). MJH and CD were supported by the University of Nottingham. We acknowledge support of the publication fee by the CSIC Open Access Publication Support Initiative through its Unit of Information Resources for Research (URICI).

## Author contributions

EP designed and performed experiments, interpreted data and wrote the manuscript.

RRT designed and performed metaphloem related experiments, interpreted data.

NR-S performed experiments and interpreted data.

FD performed LC/MS/MS and analyzed data

EK designed AtMC3 localization experiments, interpreted data.

CD performed hypoxia stress experiments, interpreted data.

RdP performed LC/MS/MS and analyzed data

DV generated AtMC3 promoter-reporter constructs

JS-L performed immunoblots

HH performed GUS stains, interpreted data

MM performed LC/MS/MS and analyzed data

MS generated AtMC3 promoter-reporter constructs

HT interpreted data and reviewed the manuscript

FVB provided materials and reviewed the manuscript.

MV designed experiments, interpreted data and reviewed the manuscript.

SM-B was in charge of hormone quantification, interpreted data and reviewed the manuscript.

MJH conceptualized hypoxia-related research, interpreted data and reviewed the manuscript.

PFH analyzed LC/MS/MS data and reviewed the manuscript.

AR-V conceptualized vascular-related research, designed experiments, interpreted data and wrote the manuscript.

NSC conceptualized the research, designed experiments, interpreted data and wrote the manuscript.

---

**Supplementary Figure 1:**
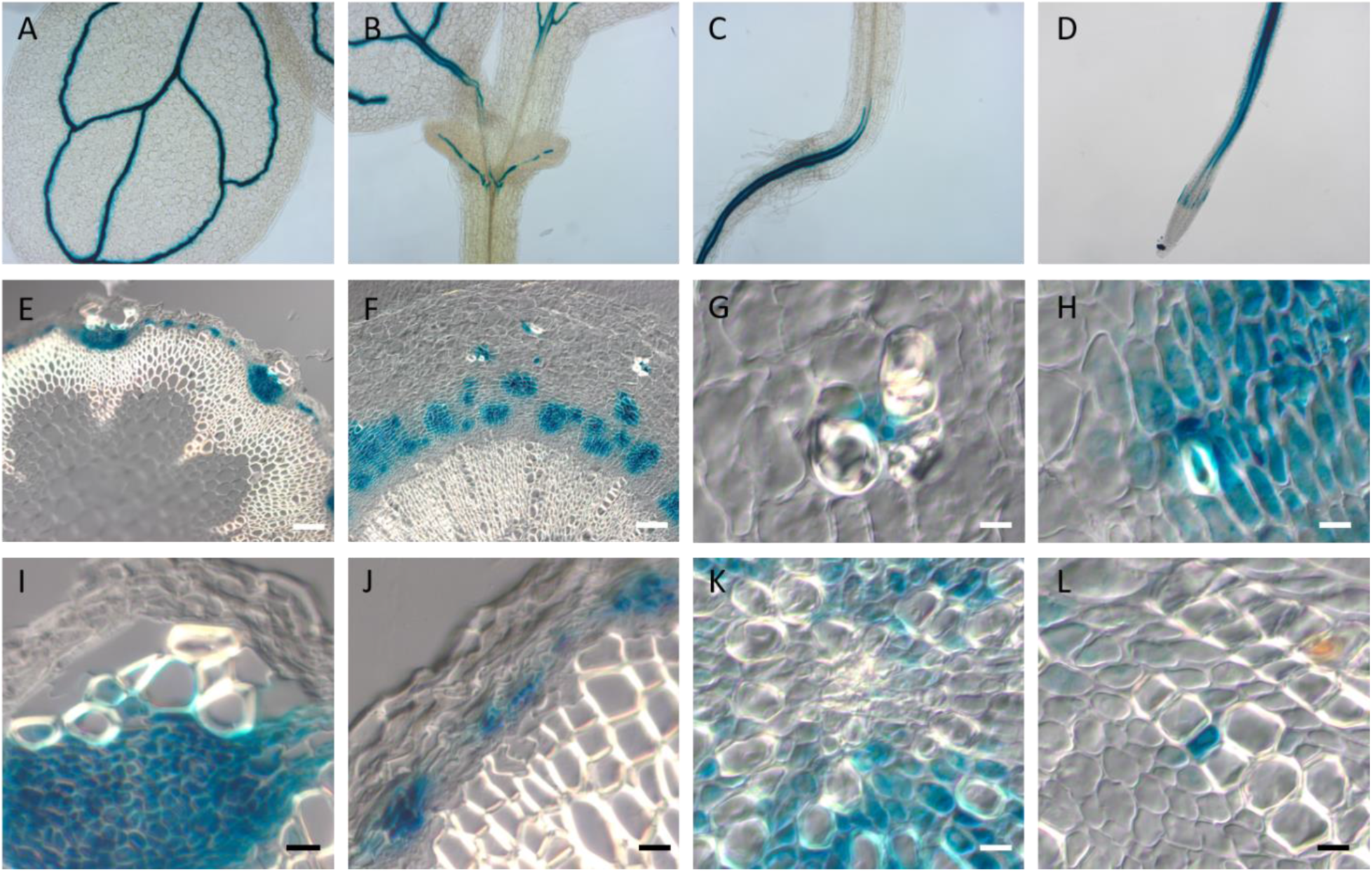
Localization of AtMC3 promoter activity in transgenic lines carrying *pAtMC3-GUS* promoter fusions. (A-D) Cotyledons (A, B), hypocotyl root junction (C) and root (D) in whole mounts of 7-days-old seedlings grown *in vitro*. (E-L) Sections from the stem (E) and hypocotyl (F-L) of 2-months-old plants grown in soil. Scale bars indicate 100 µm (E, F) or 25 µm (G-L).

**Supplementary Figure 2:**
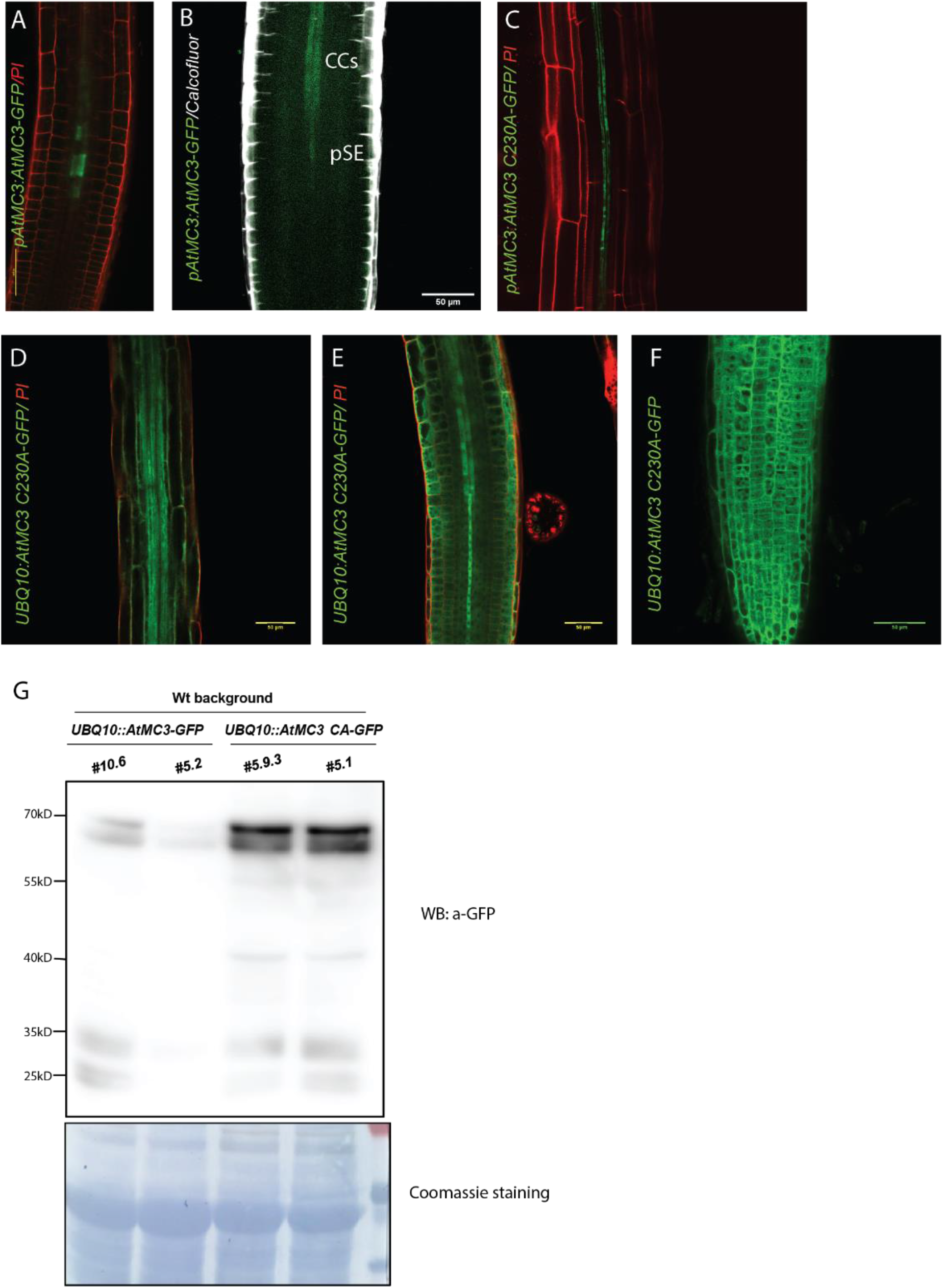
Localization of AtMC3 catalytic inactive version of protein in roots. (A-B) Expression pattern of AtMC3 protein monitored with *pAtMC3:AtMC3-GFP* in the root meristematic zone in 7-day-old seedlings. (C) Expression pattern of AtMC3 monitored with *pAtMC3:AtMC3C230A-GFP* in the roots of 7-day-old seedlings. (D-F) Expression pattern of AtMC3 monitored with *pUBQ10:AtMC3C230A-GFP* in the root of 7-day-old seedlings. (D) Differentiation zone of the root, (E) meristematic zone focusing on the vasculature, (F) meristematic zone focusing on the epidermal cells. Translational fusions with GFP signals (green) are co-visualized with propidium iodide (red) or calcofluor (white). Scale bar, 50 μm. (G) Immunoblots and respective coomassie staining for two independent lines of 7-day old seedlings carrying *pUBQ10:AtMC3-GFP* and *pUBQ10:AtMC3C230A-GFP* respectively.

**Supplementary Figure 3:**
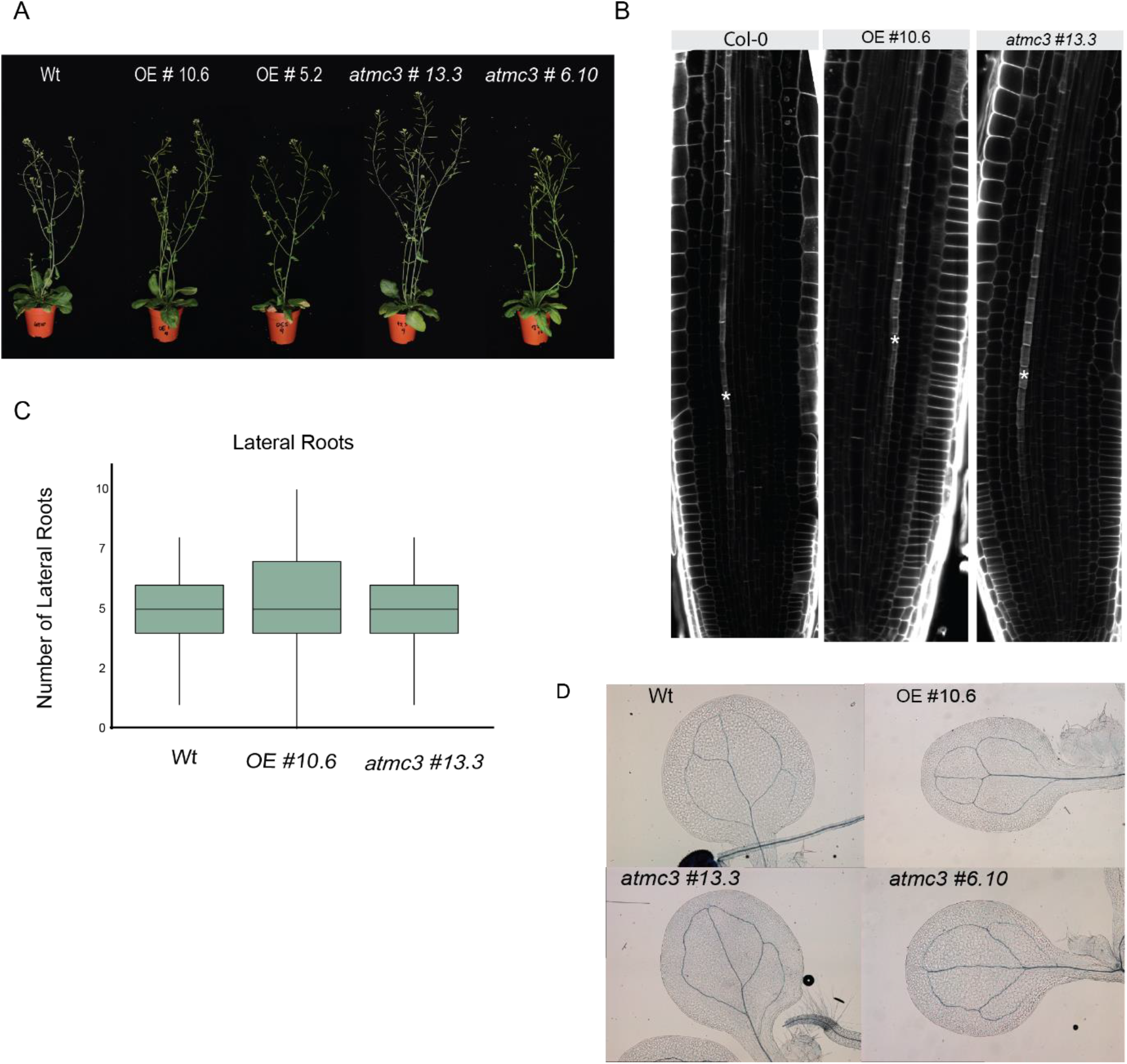
Phenotypical analysis of AtMC3 overexpressor and mutant lines. (A) Four-weeks-old representative plant phenotypes of WT, two independent AtMC3 overexpressor lines and two CRISPR *atmc3* mutant lines grown in long-day (LD) conditions. (B) Meristematic zone of the root from 6-day-old seedlings focusing on the vasculature. Visualization with Calcofluor staining. Asterisk is indicative for the differentiated PSE strand. (C) Emerged lateral roots were measured in 10-day-old seedlings grown in LD conditions for WT, overexpressor and *atmc3#13.3* lines. Three independent replicated were performed, n>30. Different letters depict significant differences within each genotype in a one-way ANOVA plus a Tukey’s HSD test. (D) Analysis of cotyledon vein pattern from 7-days-old seedlings collected from WT, overexpressor and CRISPR *atmc3* mutant plants generated.

**Supplementary Figure 4:**
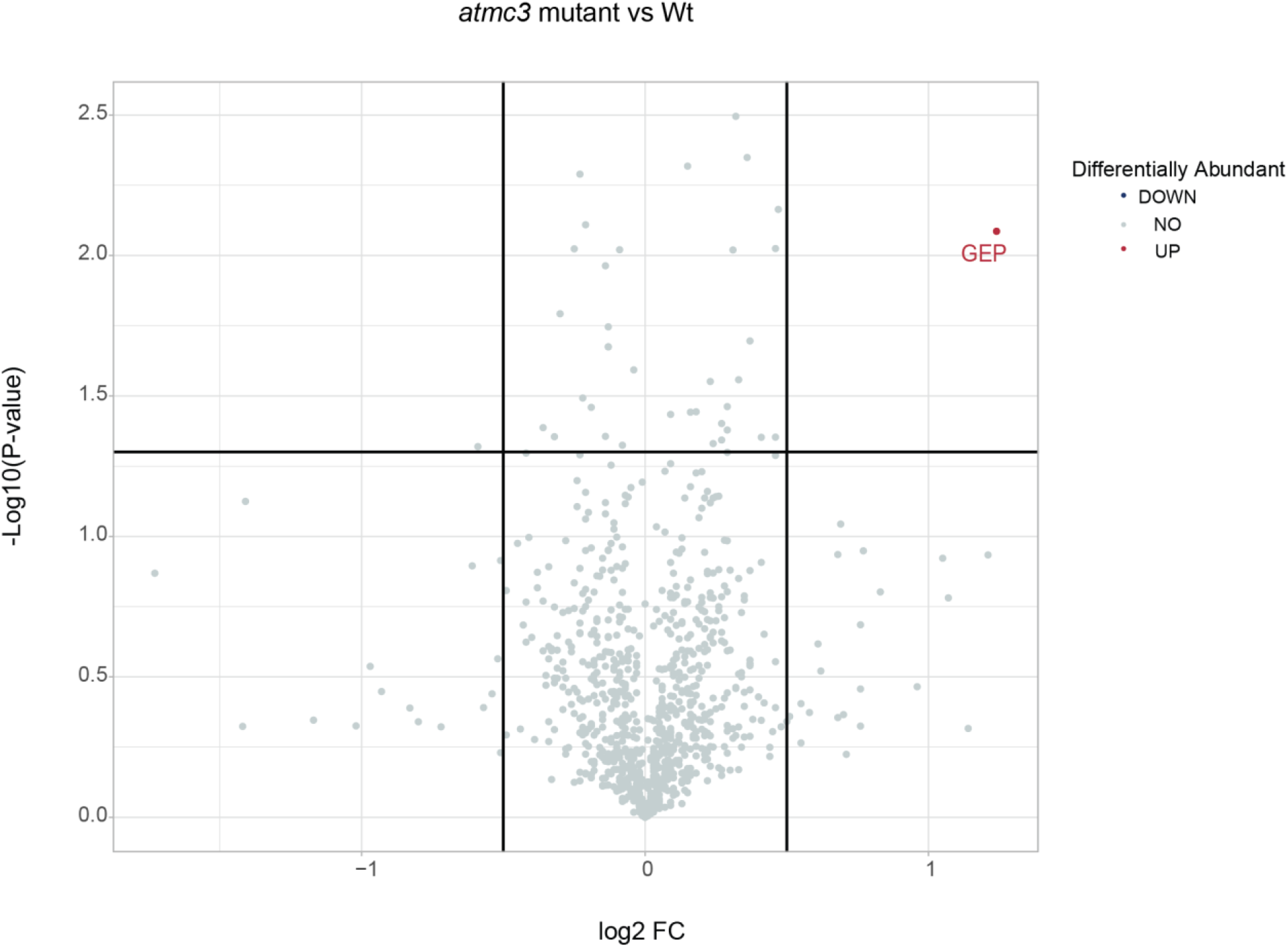
Proteomic analysis in root tissue between WT and *atmc3#13.3* mutant plants. Volcano plots of peptide abundance in WT plants compared with the *atmc3#13.3* mutant plants. N-terminal peptides with significantly reduced or increased abundance (Student′s t-test P-value<0.05 and log2 FC<–0.5 or >0.5) are highlighted with blue and red colour respectively. Results are means from four independent biological replicates.

**Supplementary Figure 5:**
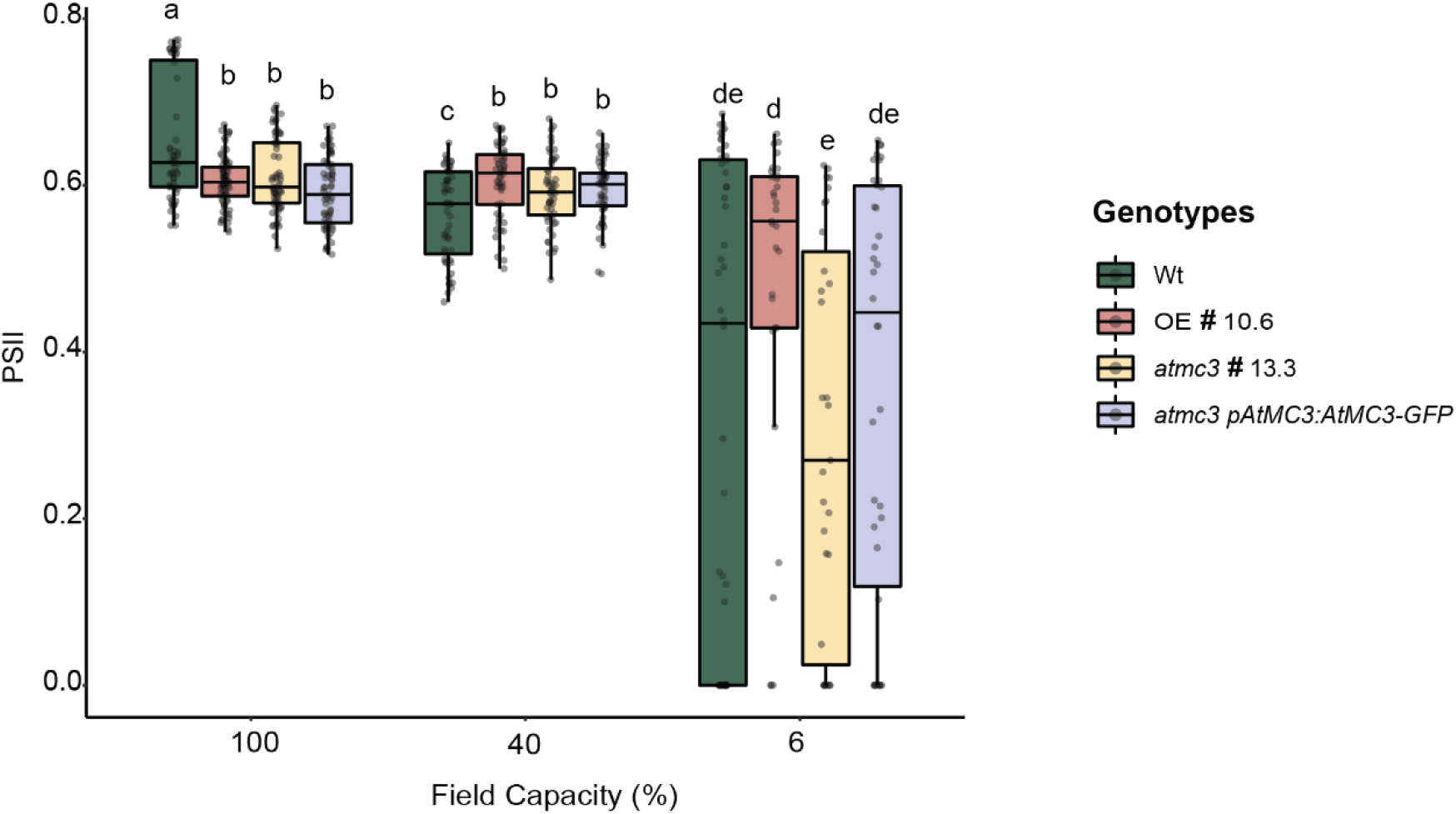
Efficiency of PSII in severe drought stress. The efficiency of PSII after exposing the plants in actinic light pulse was measured for WT, overexpressor, *atmc3* null mutant and complementation line at different percentages of field capacity. Experiments were repeated four times, n>16. Different letters depict significant differences within each genotype in a one-way ANOVA plus a Tukey’s HSD test.

**Supplementary Figure 6:**
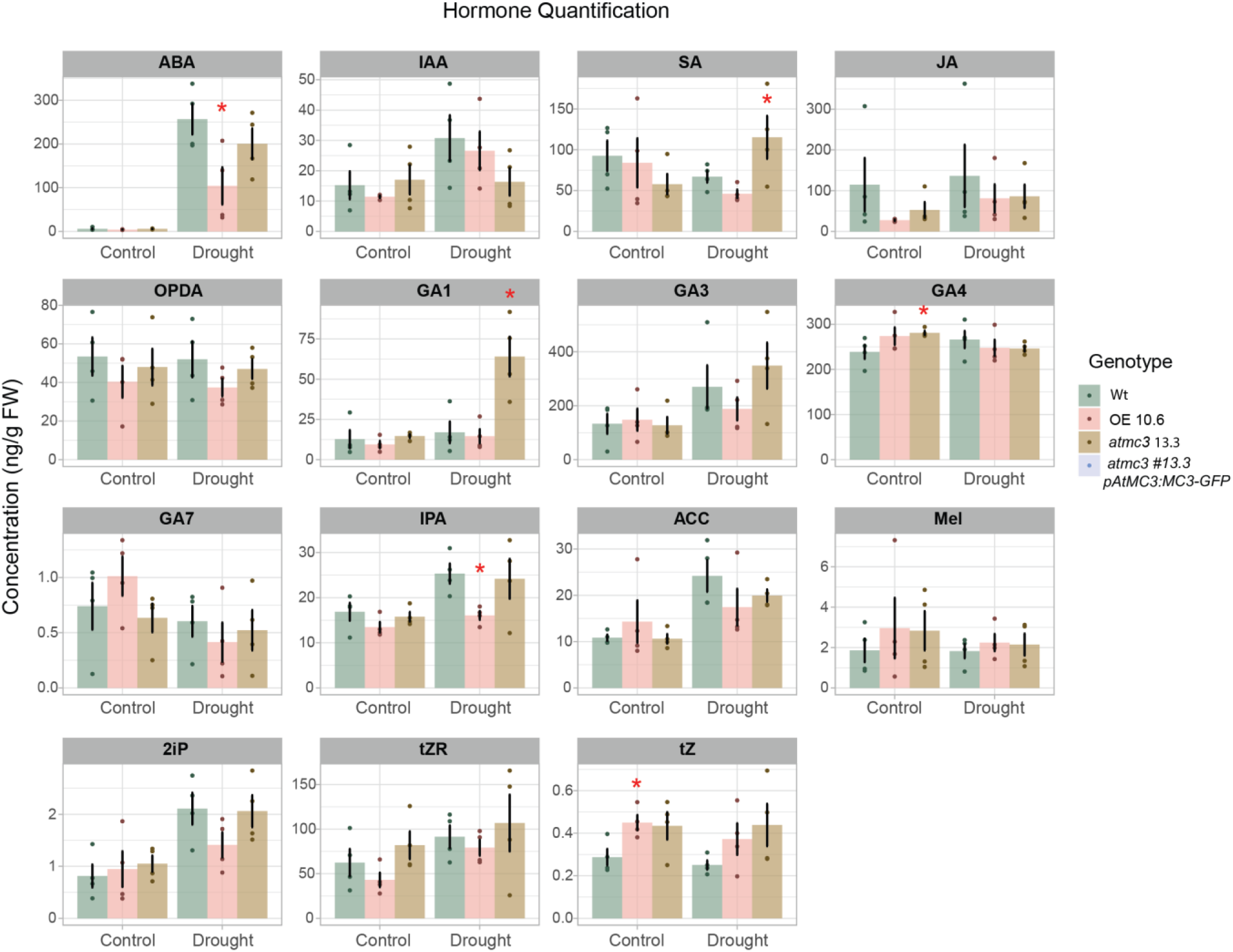
Hormonal profile in plants with different AtMC3 levels. Levels of abscisic acid (ABA), indole-3-acetic acid (IAA); the most common auxin, salicylic acid (SA), jasmonic acid (JA), 12-oxophytodienoic acid (OPDA); the precursor of JA, different types of gibberellic acid such as GA1,3,4,7, indole-3-propionic acid (IPA), 1-aminocyclopropane-1- carboxylic acid (ACC), Melatonin, 6-(γ,γ-Dimethylallylamino) purine (2iP); a type of cytokinin, trans-zeatin riboside (tZR) and trans-zeatin (tZ) were measured in WT, overexpressor and *atmc3#13.3* mutant plants. Barplots represent the results from 4 independent replicates in control (well-watered) conditions and 5 days after water withholding when plants started showing wilting symptoms. Student T-test was performed to detect significant difference in expression (* p- value< 0.05).

**Supplementary Figure 7:**
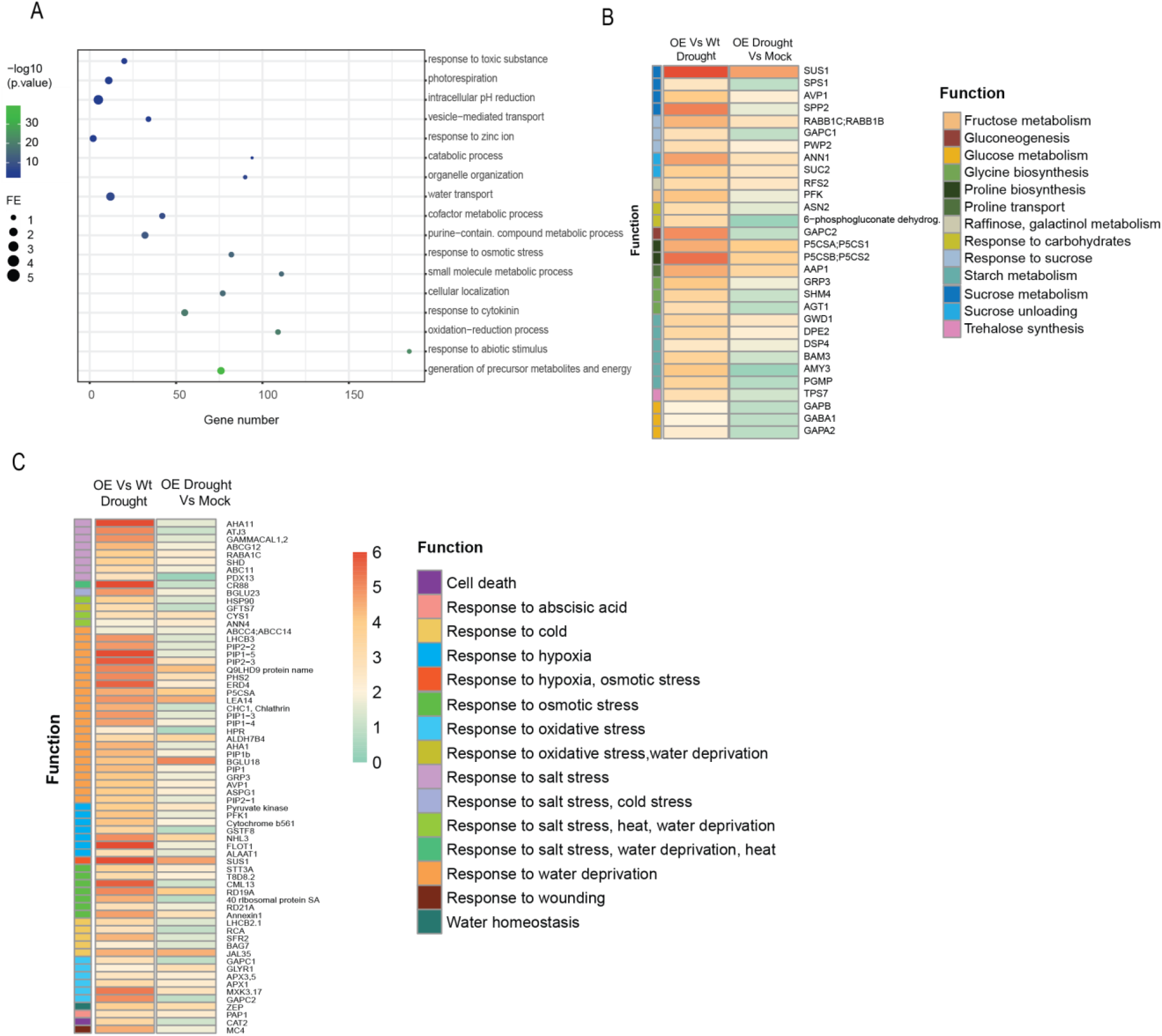
AtMC3 interactome analysis. (A) GO terms representing enriched biological processes derived from proteins co-immunoprecipitating with AtMC3 in drought stress. GO term enrichment analysis was performed on those genes that were significantly enriched in OE Vs WT plant samples under severe drought conditions (T-test, ANOVA analysis). The most representative GO provided by PLAZA^2^ was plotted along with their corresponding gene number, fold enrichment and FDR (Bonferroni Correction for multiple testing) represented as log_10_. (B, C) Heatmaps represent the log2 ratio of enriched proteins related to (B) metabolism or trafficking of osmoprotectant molecules and (C) responses to abiotic stresses in AtMC3 overexpressor /WT plants in drought conditions (left column) and AtMC3 overexpressor drought conditions / AtMC3 overexpressor mock conditions (right column). Specific colors in the squares on the left show the specific function related to the gene identified.

**Supplementary Figure 8:**
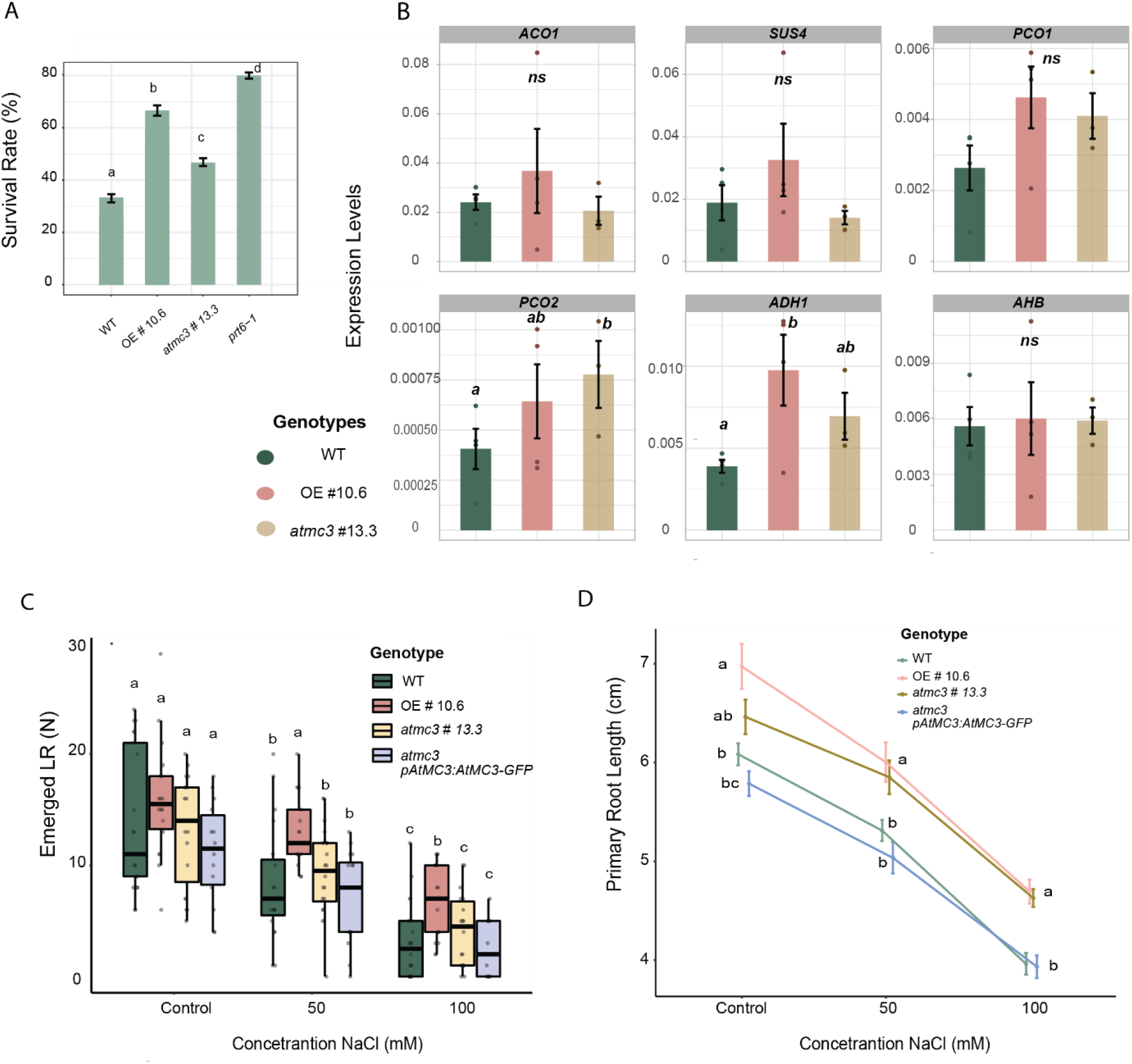
AtMC3 is involved in responses to multiple abiotic stresses. (A) Five –day-old seedlings were moved in a desiccator chamber where nitrogen was introduced at a flow rate of 4L/min in the dark, for 4hrs. Root tip survival was scored after 3 days of recovery in LD/22°C conditions in normal oxygen rate. Plant survival rates measured are indicated in the graph. Averages of three independent biological replicates (± SE), n >80. Different letters depict significant differences between the genotypes in a one-way ANOVA plus a Tukey’s HSD test. (B) Quantitative RT-PCR was performed on 7-day-old roots of seedlings grown under normoxic conditions. WT plants are compared with overexpressor of AtMC3 and *atmc3#13.3* mutant. Student T-test was performed to detect significant difference in expression between the genotypes (* p- value< 0.05). (C) Effects of salt treatment on lateral roots (LR) and (D) on primary root growth of WT, overexpressor, *atmc3#13.3* mutant and complemented plants. Different letters depict significant differences between the genotypes and the treatment (C) or between the genotypes in each treatment (D) in a one-way ANOVA plus a Tukey’s HSD test (p- value< 0.05)

Description of additional Supplementary Tables and Datasets:

**Supplementary Table 1:** Transgenic and mutant lines used in this study

**Supplementary Table 2:** Primers used for cloning strategies and genotyping

**Supplementary Table 3:** Primers used in qPCR analysis

**Supplementary Dataset S1:** List of 1072 proteins quantified Arabidopsis thaliana root samples from WT, *atmc3#13.3* crispr mutant and overexpressor line seedlings

**Supplementary Dataset S2:** List of proteins with significant changes in between overexpressor and WT root samples.

**Supplementary Dataset S3:** List of proteins with significant changes in abundance between overexpressor and *atmc3#13.3* crispr mutant root samples

**Supplementary Dataset S4:** List of proteins with significant changes in abundance between

*atmc3#13.3* crispr mutant and WT root samples.

**Supplementary Dataset S5:** Complete list of proteins that co-immunoprecipitated with AtMC3 in control and drought conditions in comparison to the WT plants under the same respective conditions.

**Supplementary Dataset S6:** Complete list of proteins that co-immunoprecipitated with AtMC3 only in drought conditions.

**Supplementary Dataset S7:** GO enrichment for Biological Process ontology in drought conditions of proteins co-immunoprecipitated with AtMC3 compared to WT.

